# PPARα-ERRα crosstalk mitigates metabolic dysfunction-associated steatotic liver disease progression

**DOI:** 10.1101/2024.10.14.618024

**Authors:** Milton Boaheng Antwi, Sander Lefere, Dorien Clarisse, Lisa Koorneef, Anneleen Heldens, Louis Onghena, Kylian Decroix, Daria Fijalkowska, Jonathan Thommis, Madeleine Hellemans, Anne Hoorens, Anja Geerts, Lindsey Devisscher, Karolien De Bosscher

## Abstract

**Background and aims:** Metabolic dysfunction-associated steatotic liver disease (MASLD) is now the most common liver disease worldwide. This study investigates how targeting two key nuclear receptors involved in hepatic energy metabolism, peroxisome proliferator-activated receptor alpha (PPARα) and estrogen-related receptor alpha (ERRα), impacts MASLD.

**Methods:** The PPARα agonist pemafibrate and/or ERRα inverse agonist C29 were administered in a short- and long-term Western diet plus fructose model, and a diabetic-background streptozotocin-Western diet model (STZ-WD). Liver morphology, histological samples, serum metabolites, RNA and protein levels were analysed and scanning electron microscopy was performed. In addition, we performed cell-based assays and immunohistochemistry and immunofluorescence stainings with light and super-resolution confocal microscopy of healthy, MASLD and MASH human livers.

**Results:** The ligand combinations’ efficacy was underscored by reduced liver steatosis in all the mouse models. Both long-term models showed improvements in body weight and liver morphology, alongside reductions in inflammation and fibrosis. Additionally, tumor formation was prevented in the STZ-WD mice model. Cell-based assays demonstrated that ERRα inhibits PPARα’s activity, likely explaining why ERRα blockage improves inflammatory and lipid metabolism gene profiles and enhances lipid-lowering effects. Complementary RNA sequencing and shotgun proteomics, combined with enrichment analysis, jointly identified downregulated serum amyloid A1/A2 as an essential component underlying the combination treatment’s effectiveness. MASLD/MASH patient livers showed reduced PPARα and increased ERRα levels supporting disrupted NR crosstalk in the hepatocyte nucleus.

**Conclusion:** Our study comprehensively supports that dual nuclear receptor targeting by simultaneously increasing PPARα and diminishing ERRα activity may represent a viable novel strategy against MASLD.

**Lay summary:** Our study using three distinct MASLD mouse models shows that a simultaneous targeting of PPARα and ERRα reduces liver fat, fibrosis, inflammation, and tumor formation. The combination treatment modifies lipid metabolism pathways, and uniquely lowers levels of serum amyloid A1/A2 in the liver.

**Impact and implications:** Our research introduces a novel therapeutic strategy against MASLD by simultaneously increasing PPARα activity while diminishing ERRα activity. With PPARα agonists already tested in phase III clinical trials, ERRα ligands/modulators need further (clinical) development to make our findings applicable to both MASLD patients and physicians.

## Introduction

Metabolic dysfunction-associated steatotic liver disease (MASLD), formerly known as non-alcoholic fatty liver disease (NAFLD), encompasses a continuous spectrum of liver disease linked to metabolic conditions such as obesity and type II diabetes mellitus.^1^ It is characterised primarily by excessive hepatic lipid accumulation termed steatosis.^2^ Currently, 30% of adults worldwide suffer from MASLD, and this number is rapidly increasing.^3^ Left untreated, MASLD can progress to an inflamed state called metabolic dysfunction-associated steatohepatitis (MASH) and further to liver fibrosis, cirrhosis, and hepatocellular carcinoma (HCC).^4^ Nuclear receptors (NRs) remain among the most relevant classes of drug targets actively investigated for MASLD.^5^ Testimony hereof is the selective thyroid hormone receptor-β agonist resmetirom, which is a recent and first FDA-approved pharmacotherapy.^6^ Our study set out to explore whether other NR family members may contribute to alleviating MASLD and provide an additional, alternative therapeutic avenue.

Peroxisome proliferator-activated receptor alpha (PPARα, NR1C1), predominantly expressed in the liver, combines nutrient sensing with anti-inflammatory activities.^7^ PPARα’s relevance has been underscored by the negative correlation between PPARα gene expression in the human liver and MASLD and MASH severity.^8^ Accordingly, hepatocyte-specific PPARα knock-out (KO) mice are more susceptible to MASLD.^9^ Pemafibrate, a potent PPARα-selective modulator, reduced liver steatosis and inflammation, lowered serum triglycerides (TG) and increased high-density lipoprotein (HDL-C) while having a good safety profile compared to conventional fibrates.^10^

Estrogen-related receptor alpha (ERRα, NR3B1) is a constitutively active orphan NR pivotal in the transcriptional control of genes relevant to mitochondrial bioenergetics and energy homeostasis.^11^ ERRα recruits the cofactor PGC1α to regulate genes encoding enzymes for fatty acid β-oxidation, the tricarboxylic acid cycle, and mitochondrial oxidative phosphorylation.^11^ Interestingly, ERRα inhibition or systemic KO of ERRα in mice protected against obesity and hepatic steatosis following chronic high-fat diet consumption and reduced hepatic de novo lipogenesis.^12,13^ Compound 29 (C29), an inverse agonist targeting ERRα, has been used chiefly for *in vitro* and preclinical studies.^14,15^

Cell-based studies and mouse models support a close functional interaction between PPARα and ERRα.^16^ Our previous study revealed crosstalk interactions between PPARα and ERRα and pinpointed ERRα as a regulator of PPARα transcriptional activity, mediated by a PGC1α-induced interaction in hepatocyte nuclei.^17^ Despite advances and independent findings linking NRs PPARα and ERRα to MASLD and MASH, the potential of a strategy targeting both NRs has not yet been investigated.^16^ Here, we assessed in three murine MASLD models whether and how simultaneous pharmacological modulation of PPARα and ERRα may interfere with MASLD disease progression. In addition, we provide data from human MASLD patients to translate our results to the clinic.

## Materials and methods

Part of the materials and methods are described in the supplementary methods.

### MASLD mouse model and pharmacological treatment

Seven-week-old C57BL/6J wild-type mice obtained from Janvier Labs, Le Genest-Saint-Isle, France, were accommodated within the Ghent University Hospital animal facility (temperature at 22°C). Mice were provided ad libitum access to standard food and water and were subjected to a 12-hour light/dark cycle. All *in vivo* experiments were ethically approved (ECD 20-87 and ECD 22-38) and conducted under the appropriate institutional and governmental regulations. Reporting adhered to the ARRIVE guidelines for animal experiments.

To assess PPARα-ERRα ligand combination efficacy *in vivo*, we used three MASLD models. In the first model, 8-week-old male C57Bl/6J mice were fed for 16 weeks with a Western diet (WD) rich in saturated fat, sucrose and cholesterol (TD.08811 + 1% cholesterol, Ssniff) and 10% fructose in the drinking water (FW). After ten weeks, mice were randomised and given vehicle (10% N,N-Dimethylacetamide, Thermo Fischer Scientific, Belgium, 20% propylene glycol, 50% polyethylene glycol 400, 20% water, Sigma-Aldrich, Belgium), PPARα agonist pemafibrate (0.1 mg/kg, 0.1 mg/kg, Bio-Connect, TE Huissen, Netherlands) and/or ERRα inverse agonist C29 (30 mg/kg, Lead Pharma, Oss, Netherlands) once via daily oral gavage for six weeks while diet feeding continued.

In the second model, C57BL/6J mice were injected subcutaneously with streptozotocin (STZ) two days after birth to destroy the pancreatic β-cells.^18^ Next, the males were fed a WD from week four to week twelve. At week six, mice were randomised and given vehicle (0.5% hydroxypropylmethylcellulose/0.1% Tween 80, Sigma-Aldrich), pemafibrate (0.1 mg/kg) and/or C29 (10 mg/kg) once daily via oral gavage for six weeks while diet feeding continued. Drug dosages aligned with prior studies.^14,15,19^

In the third model, 8-week-old male mice were fed with a WD and 10% FW for two weeks. After one week of the diet, mice were randomised and given vehicle, pemafibrate and/or C29 (30 mg/kg, Lead Pharma, Oss, Netherlands) via daily oral gavage for seven days while diet feeding continued. The animal model used, drugs, dosages and duration are summarized in Supplementary Table 1.

### Patient Samples

Human liver biopsies were obtained from healthy, MASLD and MASH patients (n=12) at the Ghent University Hospital. They were routinely processed and stained with hematoxylin-eosin (H&E) and Sirius red according to established protocols.^20^ An experienced pathologist read the biopsies, blinded to the patient’s characteristics. Histological features were scored according to the NASH clinical research network scoring system^21^. Fibrosis was evaluated using the NASH Clinical Research Network fibrosis staging system^21^. The samples were processed and subjected to dual immunofluorescence staining on the liver section for the expression and localisation of PPARα and ERRα.

### Statistical analysis

Statistical analysis was conducted using GraphPad Prism 10 (GraphPad Software Inc, USA). Quantitative image analysis was performed using Fiji (Image J, version 2.1.0) and Volocity software (Quorum Technologies In, version 7).^20^ Group differences were assessed using a one-way analysis of variance (ANOVA) with subsequent Tukey’s post-hoc testing. A two-sided P value <0.05 was deemed statistically significant. Continuous variables are expressed as mean ± standard deviation (SD).

## Results

### PPARα-ERRα ligand combination ameliorates steatohepatitis and fibrosis in a long-term MASLD mouse model

To compare the efficacy of single versus combined treatments, we employed a long-term MASLD model, administering the PPARα agonist Pemafibrate and/or ERRα inverse agonist C29. This model was designed to induce obesity with significant steatosis, inflammation, and at least a fibrosis grade 2 as established in literature.^22,23^ Hereto, mice were subjected to a 16-week regimen of a western diet (WD) and 10% fructose water (FW), including six weeks of treatment. To establish a baseline control, a subgroup was additionally sacrificed after ten weeks of WD and 10% FW diet to confirm steatosis, inflammation and fibrosis before the compound treatment was initiated (Fig.1A).

**Figure 1:**
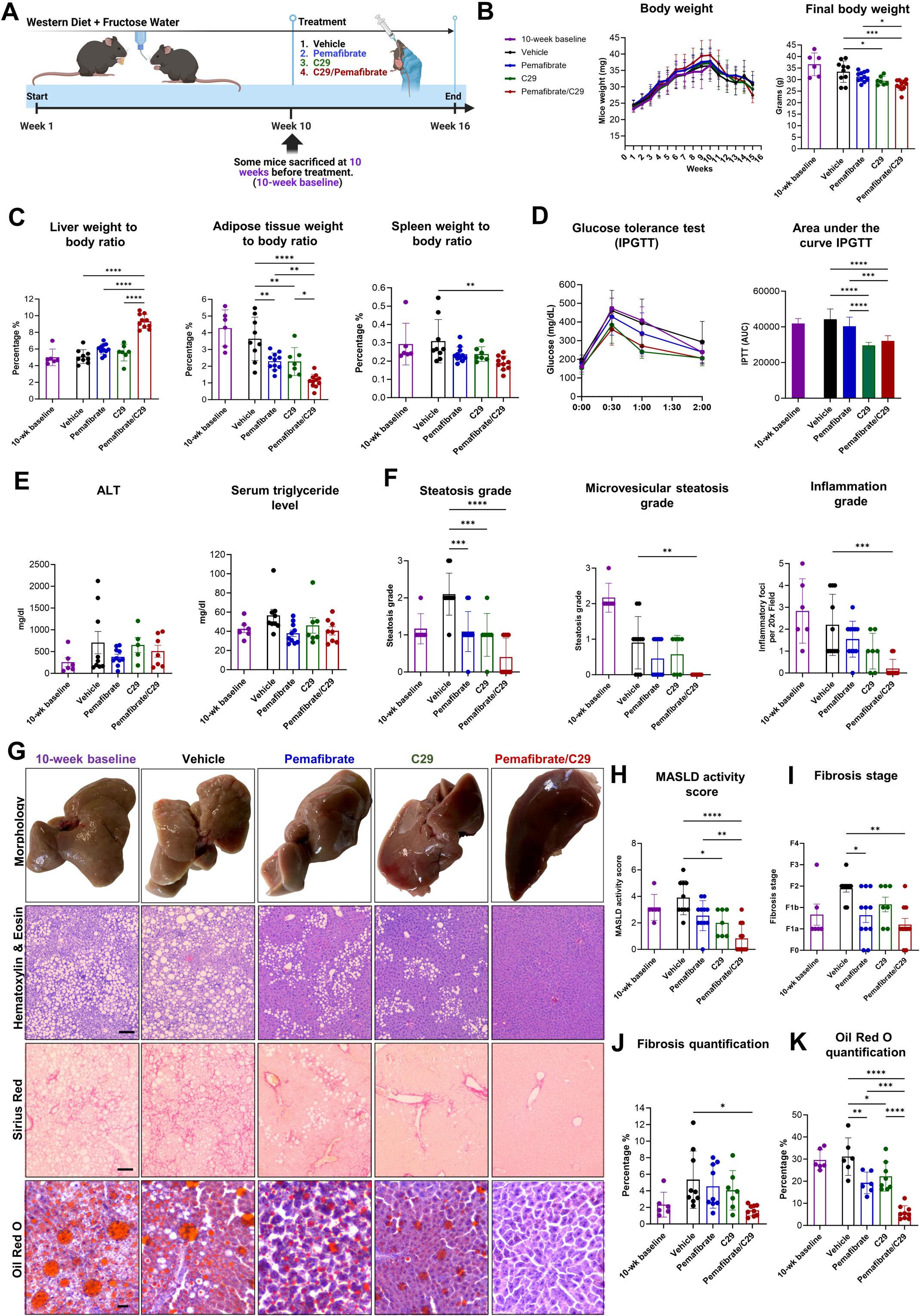
PPARα-ERRα ligand combination mitigates steatohepatitis and fibrosis in a long-term MASLD mice model. (A) 8 weeks C57Bl6 mice (7-10 mice per group) were given a western diet + 10% fructose water for 16 weeks and treated with vehicle, pemafibrate and/or C29 via daily oral gavage for 6 weeks. (B, C) Weekly body weight evolution, final body weight, liver, spleen and adipose tissue to body weight ratio. (D, E) Intraperitoneal glucose tolerance test (IPGTT) was measured, and serum from mice measured glucose, triglyceride, and alanine transaminase levels (ALT). (F-K) Steatosis, inflammation, MASLD activity score, fibrosis grade and quantification, oil red staining (ORO) results, and MASLD activity score from H&E staining. (G) Representative images of liver macroscopic morphology, H&E staining, Sirius red staining, and Oil red O staining (ORO). Representative images (10 per sample, scale bar of 100μm) were taken from Sirius red and ORO staining quantified via Fiji. All data were presented as means ± SD. The results were compared via one-way ANOVA with post-hoc testing (*: P<0.05, **P<0.01, ***P<0.001, **** P<0.0001).

Pemafibrate/C29 decreased body weight over time (Fig.1B), increased liver weight and decreased adipose tissue weight (Fig.1C), all significantly compared to single treatments and vehicle. Spleen weight was reduced, but only compared to vehicle (Fig.1C). Both C29 and the combination treatment significantly improved glucose tolerance (Fig.1D). Serum alanine transaminase (ALT) and triglyceride levels remained unaffected (Fig.1E).

Steatosis and inflammation were significantly improved comparing the ligand combination to vehicle. Only for steatosis, also single treatments showed benefit (Fig.1F). Mice treated with the ligand combination exhibited visually healthier livers (Fig.1G), in line with an improved MASLD activity score (Fig.1H). The combination treatment markedly improved fibrosis, as histopathologically staged and quantified on Sirius Red (Fig.1G,I-J), and strongly lowered lipid content, as indicated by ORO visualisation and quantification (Fig.1G,K).

Scanning electron microscopy (SEM) imaging confirmed decreased steatosis and fibrosis with the combination treatment compared to single treatments and vehicle (Fig.2A-C, Fig.S1A,B). Expression of the inflammatory genes *Tnfα and Mcp1*, as well as genes involved in fibrosis (*Col1a1, Timp1, Tgfβ*), was reduced by the ligand combination compared to vehicle (Fig.2D,E). Expression of PPARα-responsive target genes, including *Glut2, Fgf21* and *Pdk4* (glucose and lipid metabolism), *Cd36, Fabp1* (fatty acid transport), *Lpl*, and *Plin2* (lipid metabolism), were significantly and most strongly upregulated with the combination treatment (Fig.2F).

**Figure 2:**
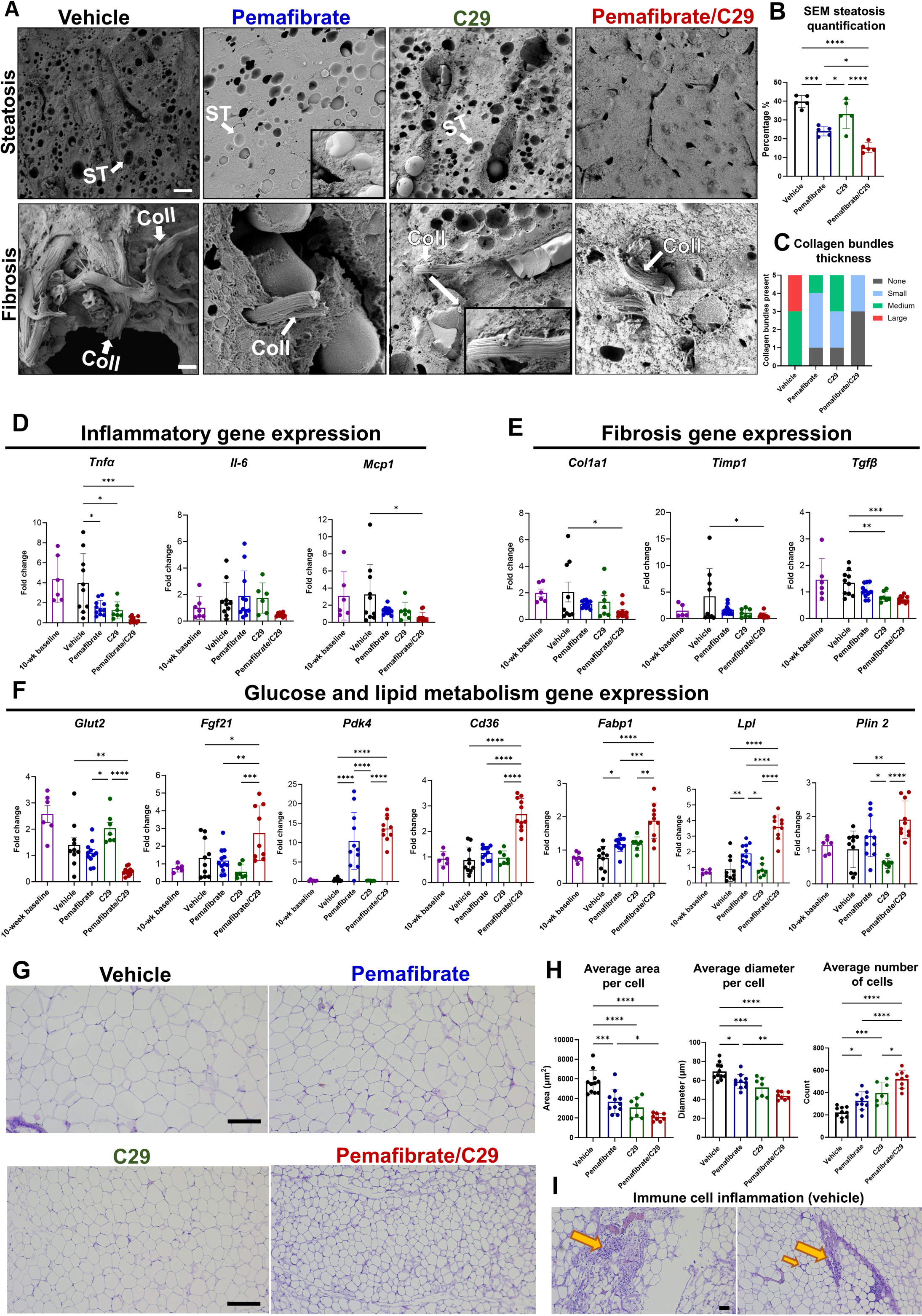
Microscopic morphology and gene expression are improved with ligand combination in long-term MASLD mice model. 8 weeks C57Bl6 mice (7-10 mice per group) were given a western diet + fructose water for 16 weeks and treated with vehicle, pemafibrate and/or C29 via oral gavage for 6 weeks. (A) Representative scanning electron microscopy images taken from the liver. (B, C) 5 images per mouse (n=4 per group) were taken to quantify steatosis (ST) and fibrosis (Coll) via Fiji (D-F) Relative gene expression levels measured by qPCR. (G-I) H&E images and quantification of adipose tissue. All data were presented as means ± SD—images scale bar 30μm for the overviews and 1μm for the high magnification images. The results were compared via one-way ANOVA with post-hoc testing (*: P<0.05, **P<0.01, ***P<0.001, **** P<0.0001).

Adipose tissue H&E staining showed a significant decrease in the average area and diameter per adipocyte with the combination treatment and an increase in the average cell number for the same surface area (Fig.2G,H). This change corresponded with a decrease in adipose tissue weight (Fig.2C). Adipose tissue immune cell infiltration was consistently present more in the vehicle than in treatment groups (Fig.2I).

### PPARα-ERRα ligand combination reduces steatohepatitis, fibrosis and tumour development in a MASLD-HCC model

No single mouse model can fully replicate all the intra- and extrahepatic characteristics of human MASLD/MASH.^24^ Therefore, we included an alternative mouse model (STZ-WD, Fig.3A) to investigate the potential impact of the PPARα-ERRα ligand combination on MASLD progression to HCC. The STZ-WD model closely mimics human MASLD pathogenesis, encompassing the sequential development of steatosis and inflammation, developing at least an F3 stage of fibrosis, tumours, and HCC on a diabetic background.^18^ While the majority of published data combines STZ with a High-fat diet (HFD) (HFD: 71% fat, 18% protein, and 11% carbohydrates), we opted for the Western diet (44.6% energy from fat, 40.7% carbohydrates (mainly sucrose) + 1% cholesterol).^18,25^ STZ-WD mice develop more significant steatosis and fibrosis than STZ-HFD mice at a shorter period.^18,25^ Here, we used a lower dose of C29 to avoid weight loss (Fig.1B), as these mice do not become obese; they rather lose weight. The treatment duration was 6 weeks to match the long-term WD plus FW model.

**Figure 3:**
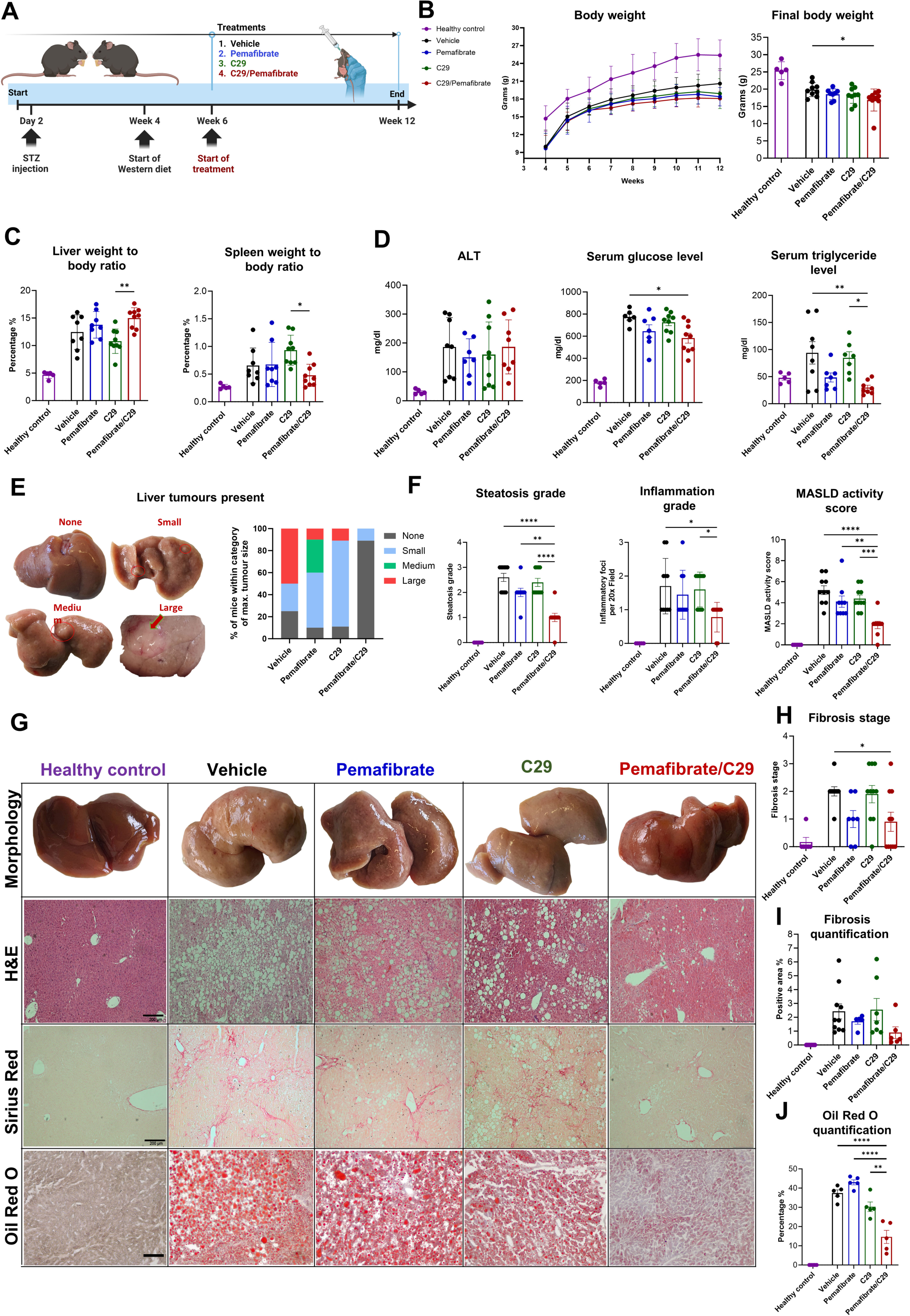
PPARα-ERRα ligand combination reduces tumours, steatohepatitis and fibrosis in the long-term diabetic Streptozotocin Western Diet mice model. (A) Mice were treated with vehicle, pemafibrate and/or C29 daily via oral gavage for 6 weeks. (B, C) Weekly body weight evolution over 12 weeks, final body weight, liver and spleen to body weight ratio. (D) Alanine transaminase (ALT), glucose, and triglyceride levels were measured. (E) Macroscopic tumour categorisation and occurrence. (F-J) Steatosis, ballooning, inflammation, MASLD activity score, fibrosis grade scoring from H&E and Sirius red staining images and Oil red O (ORO) quantification results. (G) Representative images of liver macroscopic morphology, H&E staining, Sirius red staining, and ORO staining (10 per sample). All data are presented as means ± SD—images scale bar 200μm. The results were compared via one-way ANOVA with post-hoc testing (*: P<0.05, **P<0.01, ***P<0.001, **** P<0.0001).

Gratifyingly, the results largely aligned with the long-term WD + FW mice model. The ligand combination slightly decreased body weight over time compared to vehicle. It increased liver weight and decreased spleen weight compared to C29 treatment (Fig.3B,C). Fasting glucose and triglyceride levels significantly declined with the combination treatment compared to vehicle, while ALT levels remained unaffected (Fig.3D).

The STZ-WD mice developed tumours, specifically HCC, as observed via liver morphology, H&E images, gene expression level of tumour markers (*Afp-1 and Gpc3*) and confirmed via histological staining of HCC markers (Glutamine synthase, glypican 3, and HSP70) (Fig.S2A-C). Upon combination treatment, the livers appeared healthier and displayed fewer and smaller tumours than single treatments and vehicle (Fig.2E,G). Notably, significant improvements in steatosis, inflammation, MASLD activity score, and fibrosis were again observed with pemafibrate/C29 (Fig.2F-J). As in the previous model (Fig.2D), *Tnfα* was significantly downregulated compared to vehicle, while *Il-6* and *Mcp-1* inflammatory markers only showed a decreasing trend (Fig.S2D). All treatment groups showed downregulated fibrosis markers *Col1a1* and *Timp1* (Fig.S2E). Combination treatments again decreased *Glut2* and increased the expression of PPARα target genes associated with fatty acid transport and β-oxidation, glucose and lipid metabolism consistently to the highest extent (Fig.S2F).

To explore the safety of a six-week treatment regimen with the PPARα-ERRα ligand combination separately, healthy male mice were fed a standard chow diet (Fig.S3). We observed no significant decrease in body or spleen weight (Fig.S3B). Liver weight increased with the combination treatment, while liver morphology and histology appeared normal (Fig.S3C). Serum triglyceride levels decreased upon combination treatment, whereas again, no adverse effects were observed on ALT and, importantly, on glucose levels (Fig.S3D). Interestingly, gene expression showed less outspoken crosstalk trends in a healthy setting than observed in STZ-WD and WD plus FW models (Fig.S3E). Overall, the ligand combination did not induce overt toxic side effects in healthy mice.

PPARα-ERRα ligand combination effectively decreases steatosis in a short-term MASLD model.

As a third model, mice underwent a two-week WD plus FW regimen, which induced mild steatosis, including a seven-day treatment with pemafibrate or C29 or their combination (Fig.4A). Compared to vehicle, none of the treatments altered body weight (Fig.4B), albeit pemafibrate treatments expectedly increased rodent liver weight (Fig.4C).^19^ There were no significant changes in spleen weight or glucose tolerance (Fig.4C,D). The combination treatment led to a significant decrease in fasting glucose levels compared to pemafibrate alone. All treatments reduced triglyceride levels when compared to vehicle (Fig.4D), suggesting adequate target engagement per separate ligand. Notably, the ligand combination decreased liver steatosis compared to both single treatments and vehicle already after one week of treatment (Fig.4E,F).

**Figure 4:**
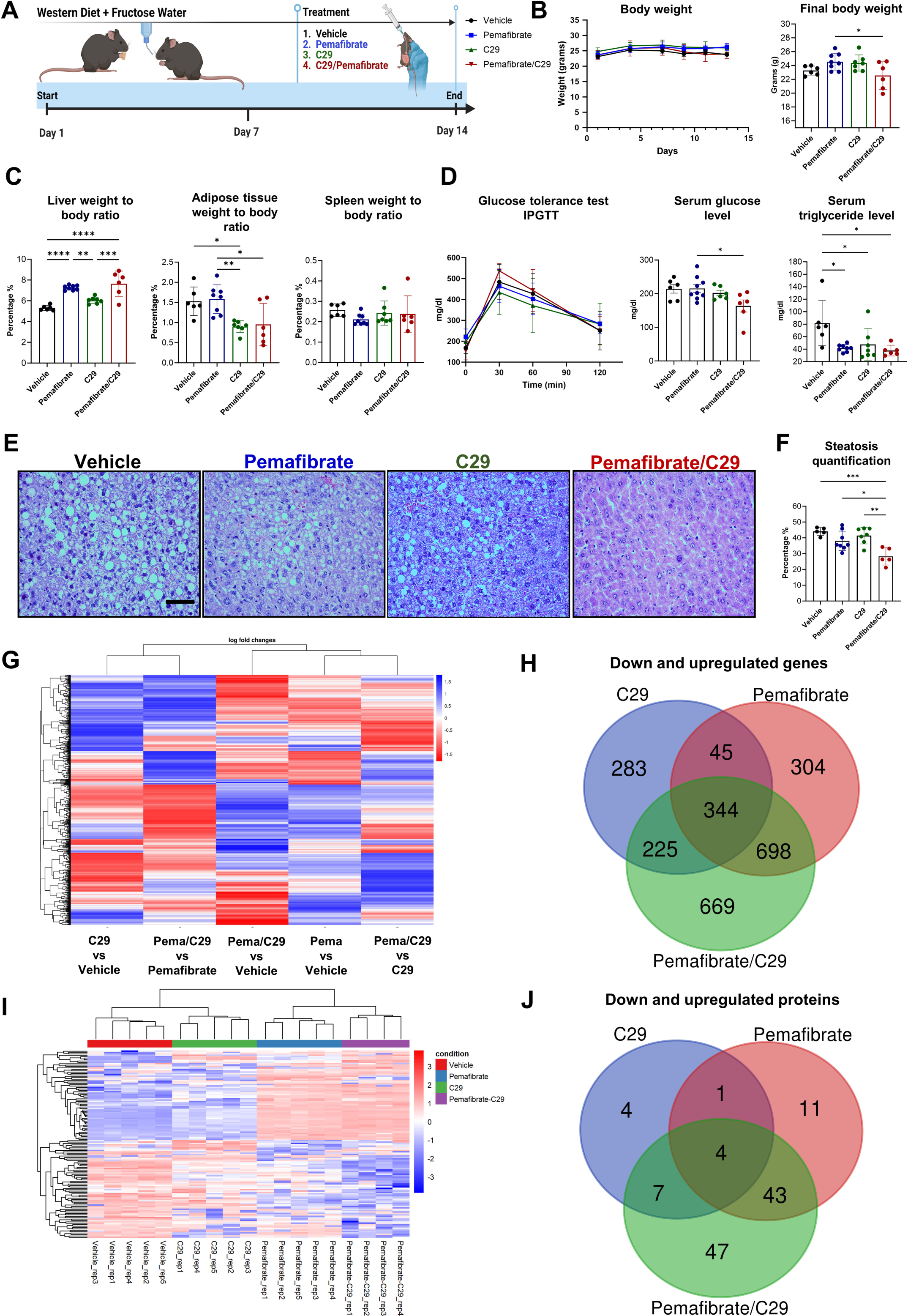
PPARα-ERRα ligand combination effectively decreases steatosis in a short-term MASLD model. (A) 8-week-old C57BL/6J mice (n=6 per group) were fed a western diet + 10% fructose water for 14 days and treated for 7 days. (B-D) Body weight evolution over two weeks, final body weight, liver, adipose tissue and spleen weight were measured. (D). Intraperitioneal glucose tolerance test (IPGTT), serum glucose and triglyceride levels were measured. (E, F) Hematoxylin and eosin staining with 5 images taken per liver section and quantified using Fiji (Image J). All data are presented as means ± SD—images scale bar 100μm. The results were compared via one-way ANOVA with post-hoc testing (*:P<0.05, **P<0.01, ***P<0.001, ****P<0.0001). (G-J). Liver samples were subjected to RNA-sequencing and MS-based shotgun proteomics (n=4-5 per group). The heatmap visually represents gene expression profiles, and the Venn diagram shows the number of shared and unique genes across treatments. Genes and proteins were filtered on p_adj_<0.05 and abs(log2FC)>=1 or >=2, respectively.

### PPARα-ERRα ligand combination selectively downregulates SAA1

To identify how pemafibrate/C29 influences the transcriptional landscape and associated protein expression, we conducted RNA sequencing and mass spectrometry (MS)-based shotgun proteomics on full liver tissue from the short-term model (Fig.4G-J and Fig.S4,S5). PCA analysis revealed four distinct clusters representing the different treatment groups (Fig.S4A, S5A). Hierarchical clustering of gene or protein expression showed that the pemafibrate/C29 combination more closely resembled the pattern of pemafibrate treatment than of the C29 treatment (Fig.4G,I). When compared to vehicle, pemafibrate/C29 yielded the largest number of differentially regulated genes and proteins (Fig.4H,J). Top upregulated genes and proteins included a.o. *Pdk4* and *Ehhadh*, which are key PPARα target genes involved in fatty acid β-oxidation (Fig.S4B,S5B) for which the combination treatment additionally increased expression compared to pemafibrate alone (Fig.S4D). Results were validated at the mRNA level via qPCR (Fig.S4D). Enrichment analysis of the genes and proteins differentially regulated by Pemafibrate/C29 vs vehicle revealed upregulated fatty acid metabolism and lipid catabolic processes (Fig.S4C,S5C).

To pinpoint the mechanisms uniquely governing the PPARα-ERRα ligand combination in the liver, we integrated RNA sequencing and proteomics data from the short-term MASLD mouse model. A drug synergy plot was calculated based on log2 fold change data of the omics analyses, indicating genes only significant upon pemafibrate/C29 versus vehicle (Fig.5A). Among the 54 unique genes/proteins identified, serum amyloid A1 (*Saa1*) stood out due to its pronounced downregulation both at the mRNA and protein level, contrasting with the unchanged or upregulated expression following single treatments. No other gene exhibited such a distinct expression pattern. GO term enrichment analysis indicated that *Saa1* is involved in the inflammatory response, specifically the acute phase response (Fig.5B). *Saa1* and its close homolog, *Saa2*, are produced by the liver during systemic inflammation and can bind to HDL particles, hereby expediting their clearance by the liver.^26^ Elevated levels of *Saa1* are linked to a higher risk of atherosclerosis and MASLD, and were shown to exacerbate hepatic steatosis^26,27^. Plotting the normalized counts for transcriptomics and proteomics of SAA1 and SAA2 confirms their substantial downregulation upon combined ligand treatment (Fig.5C,D). Additionally, we validated the expression levels of the *Saa1/2* genes in both long-term mice models and found that they were downregulated with the combination treatment (Fig.5E,F).

**Figure 5:**
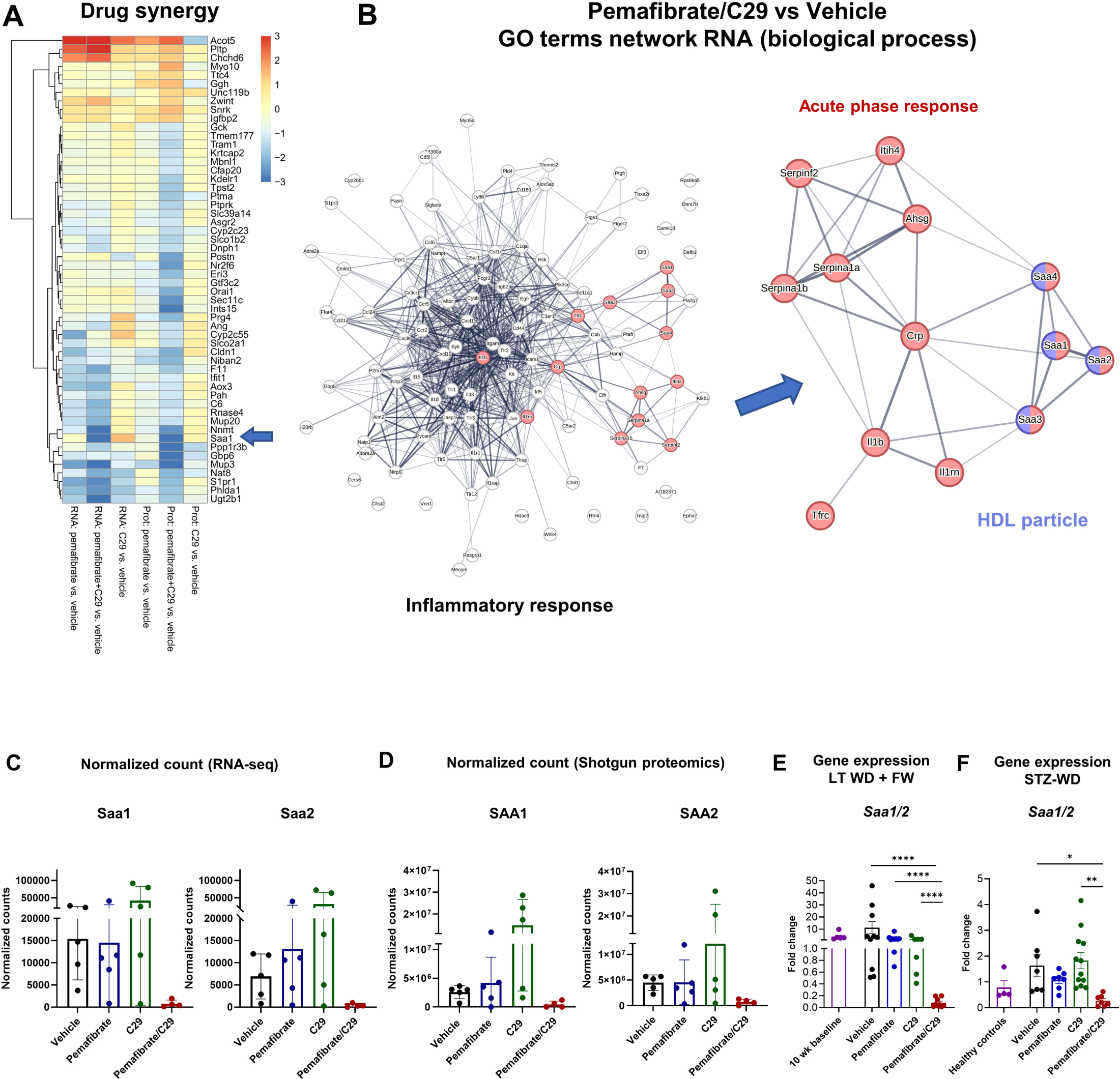
PPARα-ERRα ligand combination synergy mechanism highlighted by SAA1 downregulation. 8-week-old C57BL/6J mice were fed a western diet + 10% fructose water for 14 days, and treatment was given for 7 days. Liver samples were subjected to RNA-sequencing and shotgun mass spectroscopy (n=4-5 per group). (A) Drug synergy, calculated based on log2FC data from omics experiments, of genes significant only with pemafibrate/C29. (B). STRING-derived GO term enrichment analysis of genes differentially regulated by pemafibrate/C29 vs vehicle. (C,D) *Saa1*/2 normalized counts of RNA-seq and shotgun proteomics.

### ERRα overexpression blunts PPARα transcriptional activity

Besides our mechanistic study in whole liver, we assessed the impact of ERRα overexpression on PPARα activity in hepatocytes (Fig.S6). We used a triple PPRE-luciferase reporter in HEPG2 cells and found that ERRα suppressed GW7647-induced^28,29^, PPARα-driven transcriptional activity (Fig.S6A,C). Pemafibrate/C29 increased *Pdk4* but not *Ehhadh* mRNA expression in HEPG2 cells (Fig.S6B). To simulate steatotic conditions, cells were treated with palmitic and oleic acids before pemafibrate and/or C29, showing reduced Oil Red O staining with the combination (Fig.S6D). Overall, the data suggest that ERRα inhibition may enhance PPARα transcriptional activity in hepatocytes.

### Diminished PPARα-ERRα overlap in MASLD and MASH patients

A negative correlation between hepatic PPARα gene expression and MASH severity has been reported in mouse models and human liver samples.^8^ Because we previously observed PPARα-ERRα colocalisation in primary murine hepatocytes and in HEPG2,^17^ we additionally studied the in-situ protein expression and localisation of both NR in healthy controls (N=5) and MASLD/MASH patients (N=8) from the Ghent University Hospital (Fig.6A,B and Supplementary Table 2), who were stratified by steatohepatitis and fibrosis severity. Strikingly, PPARα-ERRα overlap was significantly reduced in patients with MASLD and MASH and patients with liver fibrosis (F2/F3) (Fig.6C-E). This was accompanied by a declining trend in total PPARα protein expression among MASLD and MASH patients, with the % of PPARα volume in the liver significantly down in case of F2/F3 fibrosis (Fig.6F). In contrast, overall ERRα protein expression seemed largely unaffected by MASLD or MASH, while F2/F3 fibrosis was associated with significantly increased ERRα protein levels (Fig.6G). Finally, we assessed potential changes in receptor subcellular localisations (Fig.S7). While the relative PPARα protein signals inside the hepatocyte nucleus were not significantly different (Fig.S7A), a downward trend was apparent outside of the nucleus, especially in patients with F2/F3 fibrosis (Fig.S7B,E). ERRα protein levels, both inside and outside hepatocytes, did not differ between patients with MASLD/MASH (Fig.S7C,D). However, outside the nucleus, ERRα signals were increased in patients with F2/F3 fibrosis (Fig.S7D,F).

**Figure 6:**
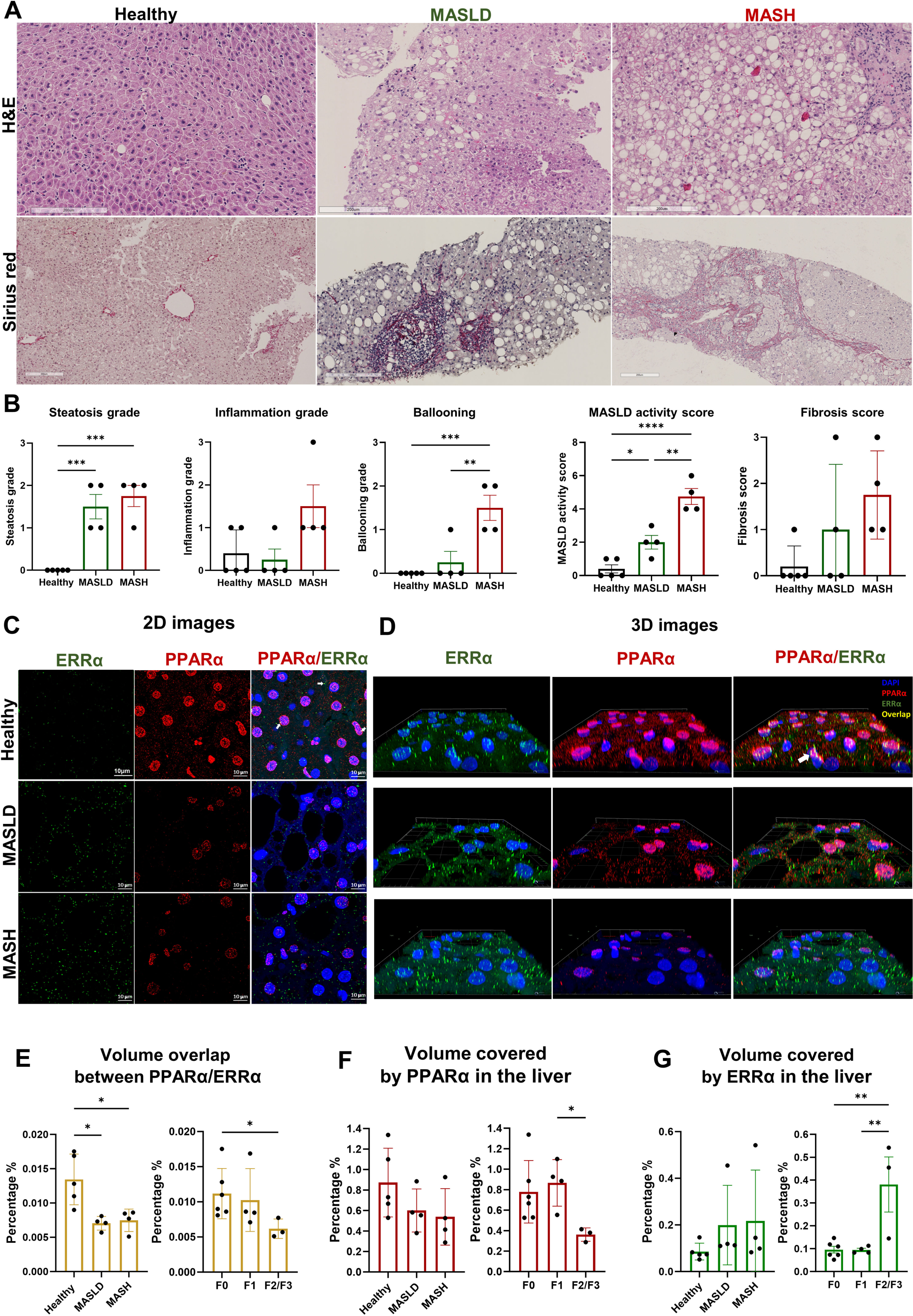
PPARα-ERRα receptor expression in healthy, MASLD and MASH patients. Liver biopsies from Ghent University Hospital were processed for double immunostaining from healthy, MASLD and MASH patients. (A) Representative H&E images and fibrosis images from liver biopsies. (B) Patient samples graded scores of steatosis, inflammation, ballooning, MASLD activity and fibrosis. (C) Representative 2D images of PPARα (red) and ERRα (green) against nuclei (blue) in the liver. (D) Magnified 3D image from healthy control showing the overlap between PPARα (turquoise), ERRα (green) and overlap between both receptors (yellow). (E-G) 6 images (approximately 60 images per Z-stack) were taken per liver section with an airy scan LSM 880 super-resolution microscope, and the endogenous protein volumes (3D images) were quantified using Volocity. We added white arrows to point to some sections of overlap. All data are presented as means ± SD—images scale bar 10μm. The results were compared via one-way ANOVA with post-hoc testing (*: P<0.05, **P<0.01, ***P <0.001, **** P<0.0001).

## DISCUSSION

PPARα, being highly expressed in the liver, regulates lipid metabolism by enhancing fatty acid β-oxidation and decreasing lipogenesis, with pemafibrate being a newer-generation PPARα-selective agonist.^19,28^ ERRα, an orphan nuclear receptor with a putative ligand binding pocket almost completely occupied by amino acid side chains,^30^ only recently gained attention as a drug target against MASLD/MASH.^15^ Diminishing ERRα’s activity via the inverse agonist C29 has been reported to ameliorate MASLD and MASH.^11,12,31^ Our investigation into PPARα-ERRα ligand combinations, encompassing three murine models and a liver cell line, supports enhanced therapeutic benefit in MASLD upon simultaneously activating PPARα while inhibiting ERRα via systemic treatment.

In our in vivo study, we utilised male mice due to their heightened susceptibility and severity to developing MASLD compared to female mice.^32^ This allowed us to comprehensively assess the efficacy of ligand combinations versus single treatments. In both long-term models (Fig. 1-3), the most striking features of the ligand combination were significantly reduced steatosis, inflammation and fibrosis, hereby outperforming the effects of single ligand treatments.

According to the literature, both pemafibrate and C29 individually reduce body and adipose tissue weight in mice, which our study confirmed.^12,14^ However, in the WD plus FW model, pemafibrate/C29 further decreased body and adipose tissue weight (Fig.1), due to a decreased adipocyte surface area and diameter per cell. Less immune cell infiltration compared to vehicle treatment was consistent with a decreased inflammatory gene profile (Fig.2). Moreover, pemafibrate and C29, when administered separately, both normalise triglyceride and glucose levels in MASLD.^10,11^ A 2020 study by Sasaki *et al*. showed that pemafibrate improved MASLD in an STZ-high-fat diet mice model but did not reduce hepatic triglyceride content.^33^ Here, using the STZ-WD mice model instead and adding C29 to pemafibrate effectively decreased liver hepatic triglyceride content (Fig.3). This was consistent with visually reduced lipid buildup with the ligand combination via ORO staining in HEPG2 cells.

Another striking result was the prevention of HCC development by the combination treatment in the STZ-WD model. In line, inhibition of ERRα as a single agent strategy is under investigation for several cancers, including breast and endometrial cancers.^34,35^ Adding a PPARα agonist to the mix may further open potential gateways toward HCC pharmacotherapy, especially in metabolic dysfunction.

Pemafibrate studies in mice have shown that liver weight/size increases in correlation with dose; the higher the dose, the greater the enlargement^19^. As inhibition of ERRα causes increased PPARα activity, this may explain the observed liver enlargement with the combination treatment in both long-term models. Spleen enlargement is commonly associated with MASLD, as well as liver and portal fibrosis.^36^ Unlike the individual ligands, the ligand combination effectively reduced spleen weight, apparent in the long-term WD + FW mice model (Fig.1). Our parallel study and findings in healthy mice confirm the safety and tolerability of PPARα-ERRα ligand combinations. While expectedly displaying a surge in fatty acid oxidation, the ligand combination did not provoke disruptive alterations of physiological cellular processes.

Even though the beneficial effects of the ligand combination over individual treatments were most outspoken in the long-term MASLD models, notable effects in the short-term model justified a follow-up to find a mechanistic basis (Fig.4). Complementary techniques, including RNA sequencing and MS-based shotgun proteomics of whole liver as well as qPCR and reporter gene assays in HepG2 cells helped understand how the ligand combination affects gene expression and regulates pathways. Integrated RNA sequencing and proteomics data revealed drug synergy with the PPARα-ERRα ligand combination (Fig.5). Saa1/2, expressed in hepatocytes, was significantly downregulated at both mRNA and protein levels with the combination treatment, which may be one explanation for why the drug combination is so effective. SAA1 is elevated during inflammation, MASLD/MASH and cancer in both mice and humans.^27,37,38^ Studies support that inhibiting SAA1 alleviates obesity and insulin resistance.^39,40^ In a study in which mice were fed a high-fat diet (HFD) for 16 weeks, knocking out SAA1 alleviated hepatic steatosis and inflammation.^27^ Interestingly, HFD-induced hepatic overexpressed SAA1 aggravates liver inflammation in MASLD, whilst blocking SAA1 alleviates it.^38^ These findings highlight the role of Saa1/2 in toxic lipid accumulation and pathogenic inflammation during MASLD progression. Our long-term mouse models confirmed Saa1/2 downregulation with the combination treatment, a distinct effect not observed with the single treatments.

In our study, inhibiting ERRα consistently increased PPARα activity in the diseased mouse livers, resulting in the upregulation of PPARα target genes such as e.g. *Pdk4* and *Ehhadh* with the latter examples encoding classical peroxisomal β-oxidation enzymes.^41^ Confirming the gene expression data and in support of a direct transcriptional crosstalk mechanism, ERRα overexpression strongly repressed PPARα-driven gene expression at an isolated PPRE-dependent promoter (Supp. Fig. 6). This aligns with our recent study, where inhibiting ERRα pharmacologically reduces its interaction with ligand-activated PPARα and results in increased mRNA expression of key PPARα target genes, indicating that ERRα may act as a transcriptional repressor^17^. More evidence in support of PPARα and ERRα being molecularly intertwined was found in other model systems.^12,17,31,42^ Although combining both NR ligands may target other liver cells and pathways involved in fibrogenesis and inflammatory responses, as observed with different drug combinations in MASLD, hepatocytes seem a most likely target given their high NR content, mirroring of lipid reduction, and at least the *PDK4* gene expression profile in the HEPG2 model.^43^

Augmenting insights, patient data from Ghent University Hospital provided a clinical lens across disease states (Fig.6). The observed downregulation of PPARα protein aligns with previous studies that linked a lowered hepatic PPARα expression with disease severity in MASH patients.^8^ Although ERRα is known to be transcriptionally upregulated in HFD-fed mice,^11^ changes in ERRα protein expression and subcellular localisation in MASLD patients have not yet been reported. While our study showed no apparent differences in ERRα levels in MASLD and MASH patients, ERRα levels were elevated in patients with F2/F3 fibrosis. A notable observation was the decrease in overlap between both NRs in MASLD, MASH and patients with severe fibrosis as compared to healthy individuals. The diminished PPARα levels coinciding with elevated ERRα levels may form the basis for a disrupted NR crosstalk mechanism during MASLD progression. One implication of this finding is that in concert with trying to enhance PPARα, ERRα may even be more accessible to efficient targeting in the liver because of its enhanced levels. It should still be considered that the function of human and mouse PPARα isoforms has species-dependent features,^7^ potentially explaining why solely triggering PPARα significantly improves murine MASLD, while its effects in clinical studies are minor. Enhancing PPARα activation through repression of ERRα might be an avenue to strengthen clinical effects.

Our findings collectively suggest that the PPARα-ERRα ligand combination holds therapeutic promise to tackle MASLD progression. The consistency of positive outcomes across models and the translational insights from human data emphasise the need to further explore the potential clinical relevance of this approach. Future studies focusing on integrating (spatial)omics technologies may further refine our understanding of PPARα-ERRα crosstalk to inhibit and ameliorate MASLD progression. A future translational path should help fine-tune therapeutic strategies aimed at enhancing PPARα activity while mitigating ERRα’s counteractive effects.

## Author contributions

Conception and design: KDB, AG, LD, SL, MA; Supervision: KDB, AG, LD, SL; Animal experiments and data analysis: MA, SL, AHe, LO; In vitro experiments and analysis: MA, KD, MH, JT; Human sample collection and analysis: MA, SL, AG and AHo; RNA sequencing data analysis: JT, LK, DC and MA; Mass spectrometry analysis: DF, DC, MA; Critical revision for important intellectual content: KDB, AG, LD, SL, LK, DC; Writing original draft: MA; Writing review and editing: KDB, LD, AG, SL, DC, MA; All authors reviewed and approved the manuscript.

## Acknowledgements

We warmly thank Petra Van Wassenhove, Inge Van Colen, and Els Van Deynse for the excellent technical support, for providing biochips, and all members of our labs and collaborating scientists for helpful discussions. We also thank the VIB Nucleomics and Proteomics Core for adequate processing and assistance with the analysis of our samples and the VIB Ghent Bioimaging Core scientists for providing technical support during confocal microscopy and image analysis. We would also like to thank Lead Pharma for providing us with the C29 compound and handling details.

