## Supplementary figures, tables, and methods for "PPARα-ERRα crosstalk mitigates metabolic dysfunction-associated steatotic liver disease progression"

#### Contents

Supplementary figures 1-7 page 2

Supplementary figure legend page 9

Supplementary tables page 12

Supplementary methods page 14

Supplementary method references page 27

**Figure S1**

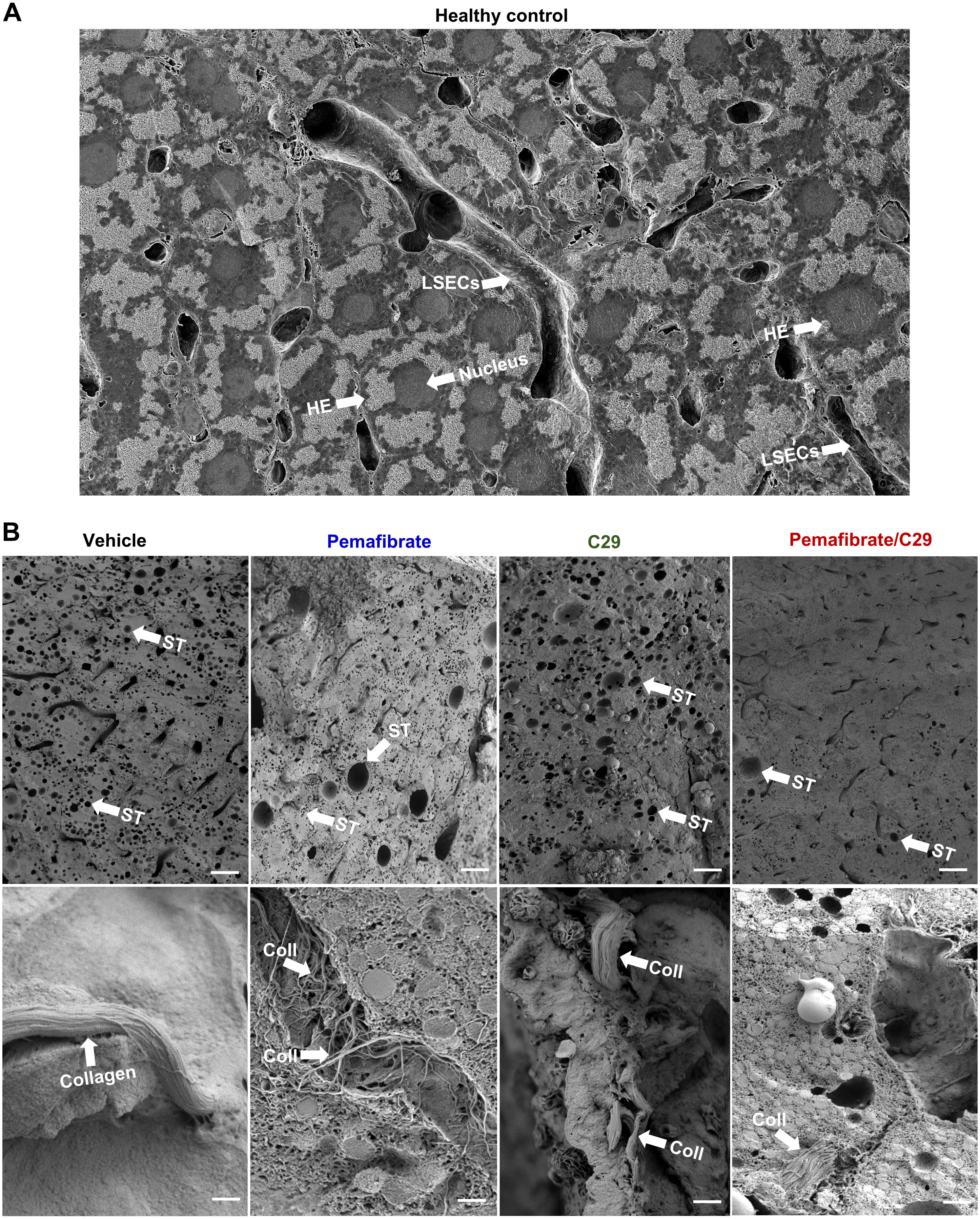
A
Healthy control
LSECs
HE
Nucleus
HE
LSECs
B
Vehicle
Pemafibrate
C29
Pemafibrate/C29
ST
ST
ST
ST
ST
ST
ST
ST
Collagen
Coll
Coll
Coll
Coll
Coll

#### Figure S2

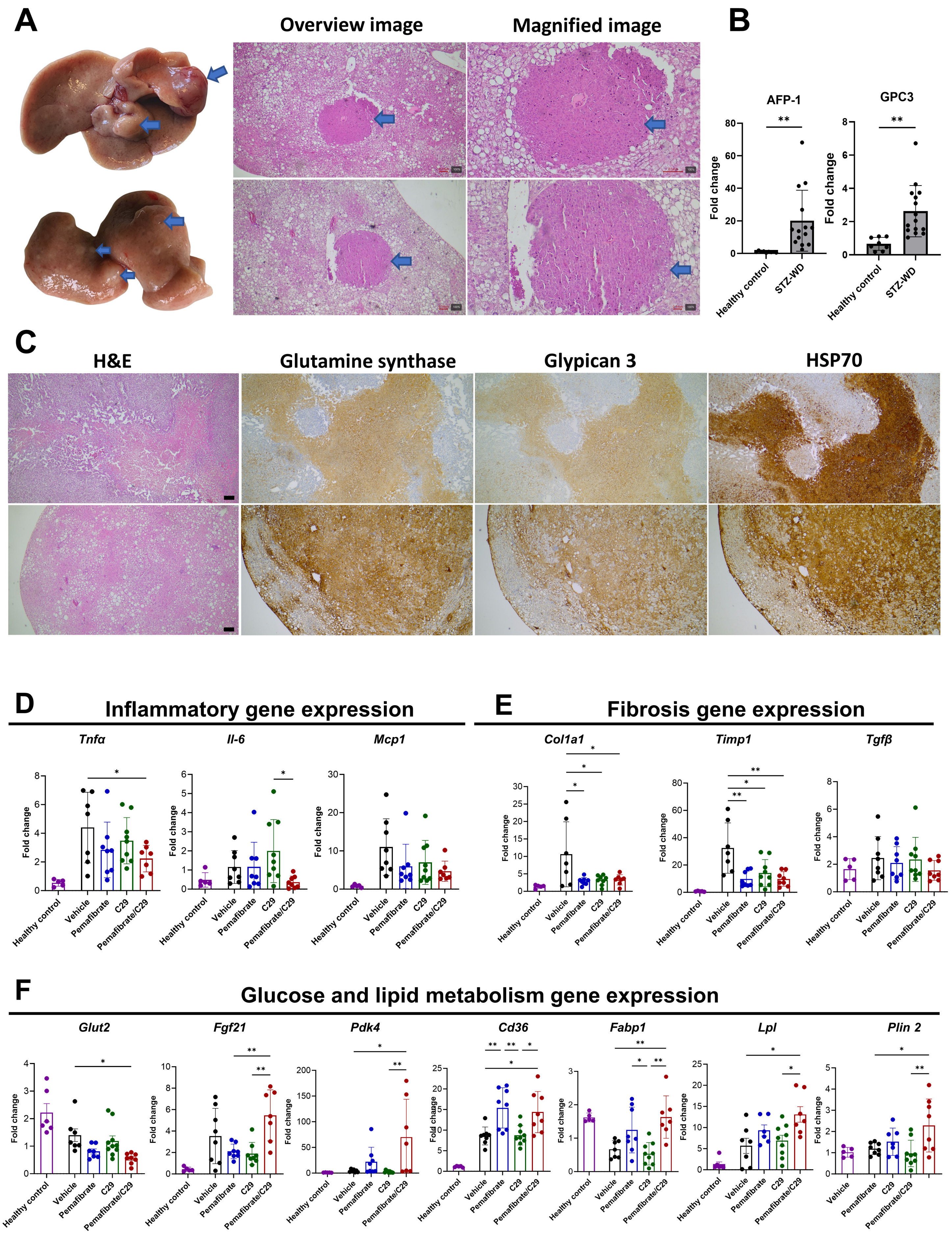
A
Overview image
Magnified image
B
AFP-1
GPC3
Fold change
Healthy control
STZ-WD
**
C
H&E
Glutamine synthase
Glypican 3
HSP70
D
Inflammatory gene expression
Tnfα
Il-6
Mcp1
E
Fibrosis gene expression
Col1a1
Timp1
Tgfβ
F
Glucose and lipid metabolism gene expression
Glut2
Fgf21
Pdk4
Cd36
Fabp1
Lpl
Plin 2
Fold change
Healthy control
Vehicle
Pemafibrate
C29
Pemafibrate/C29

#### Figure S3

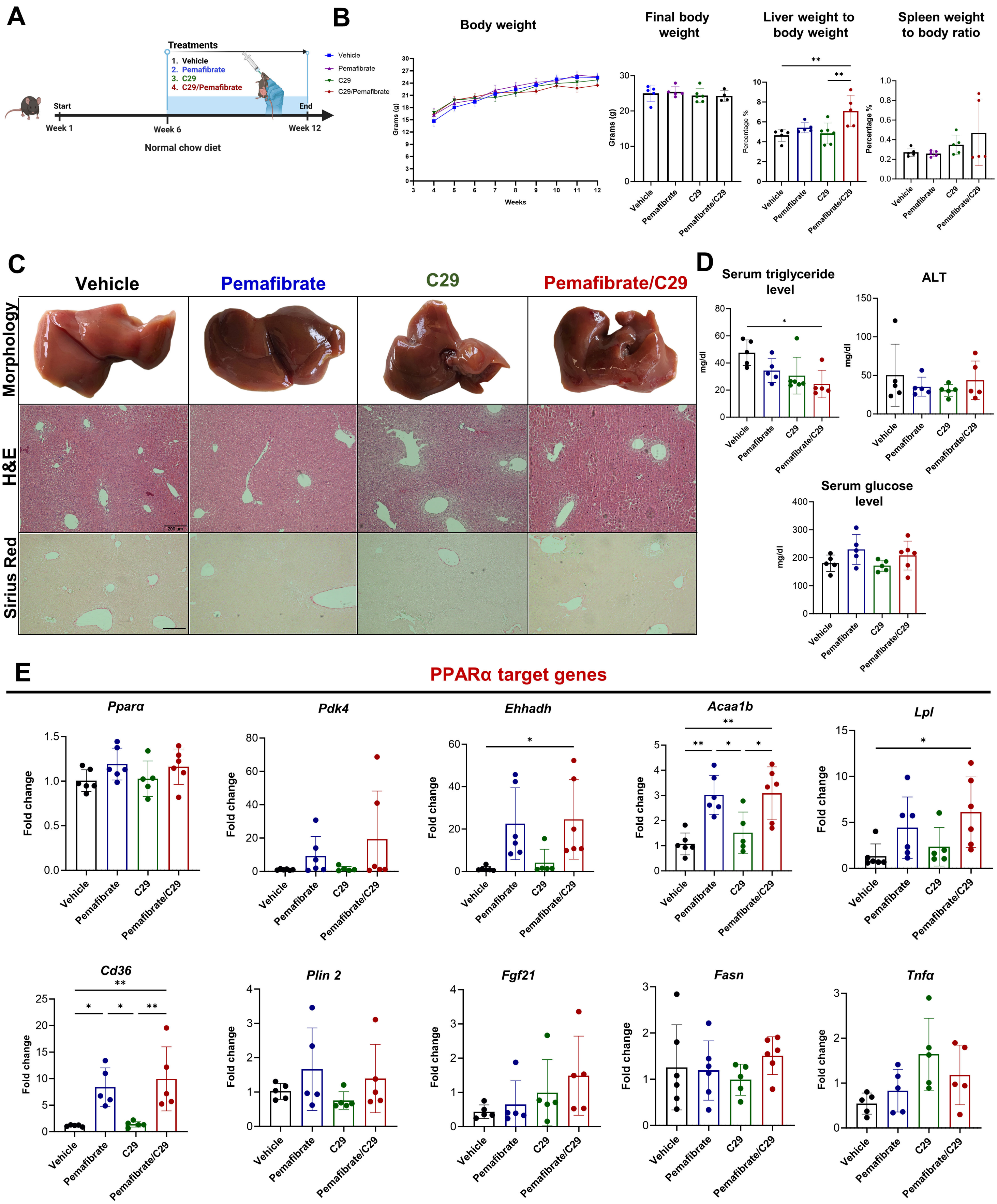
A
Treatments
1. Vehicle
2. Pemafibrate
3. C29
4. C29/Pemafibrate
Start
End
Week 1
Week 6
Week 12
Normal chow diet
B
Body weight
Vehicle
Pemafibrate
C29
C29/Pemafibrate
Grams (g)
Weeks
Final body weight
Grams (g)
Liver weight to body weight
Percentage %
**
**
Spleen weight to body ratio
Percentage %
C
Vehicle
Pemafibrate
C29
Pemafibrate/C29
Morphology
H&E
Sirius Red
200 μm
D
Serum triglyceride level
mg/dl
*
ALT
mg/dl
Serum glucose level
mg/dl
E
PPARα target genes
Pparα
Pdk4
Ehhadh
Acaa1b
Lpl
Cd36
Plin 2
Fgf21
Fasn
Tnfα
Fold change
Vehicle
Pemafibrate
C29
Pemafibrate/C29

#### Figure. S4

##### RNA-sequencing whole liver short-term MASLD mice model

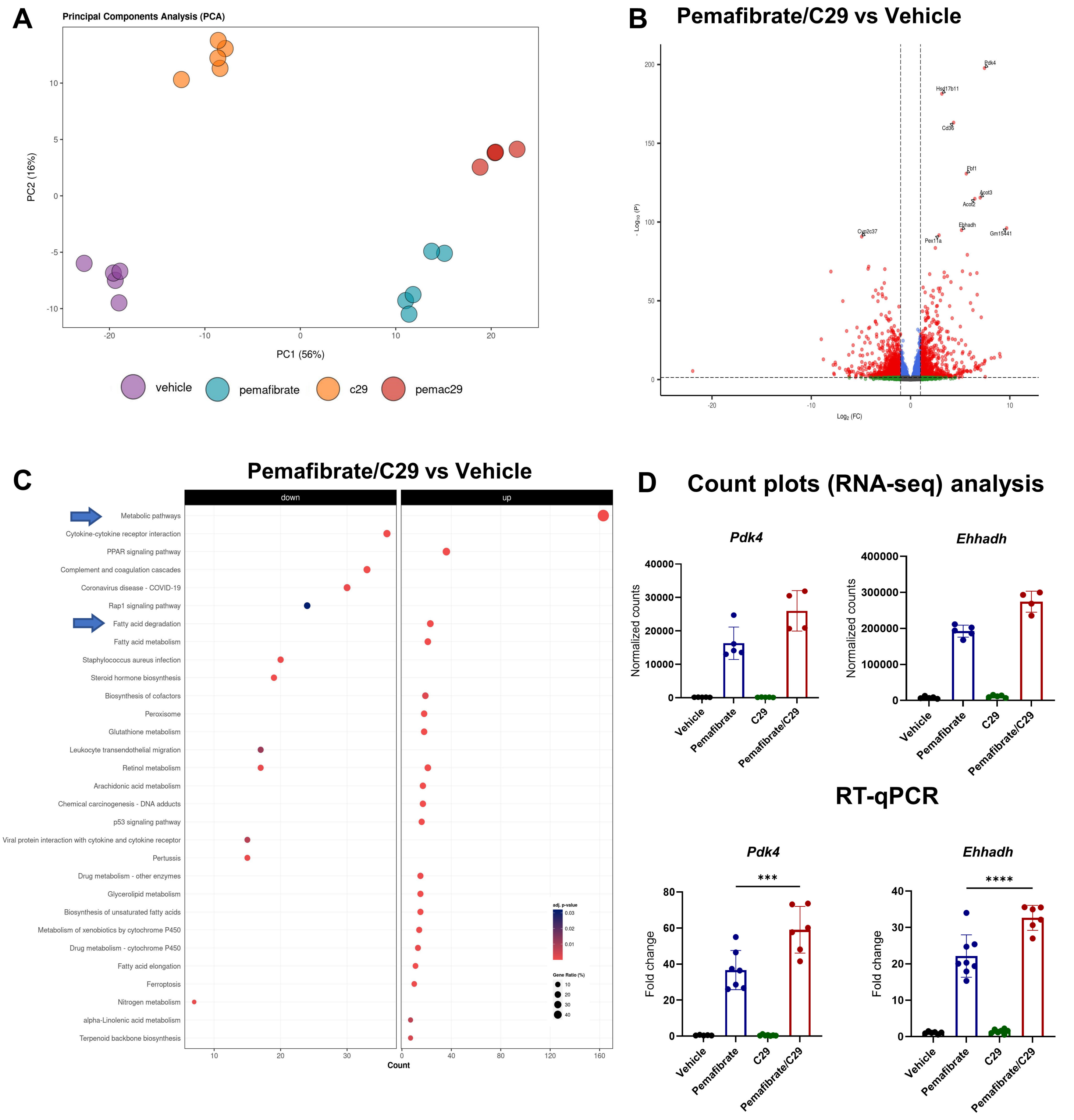

**Fig. S5**

#### Shotgun proteomics whole liver short-term MASLD mice model

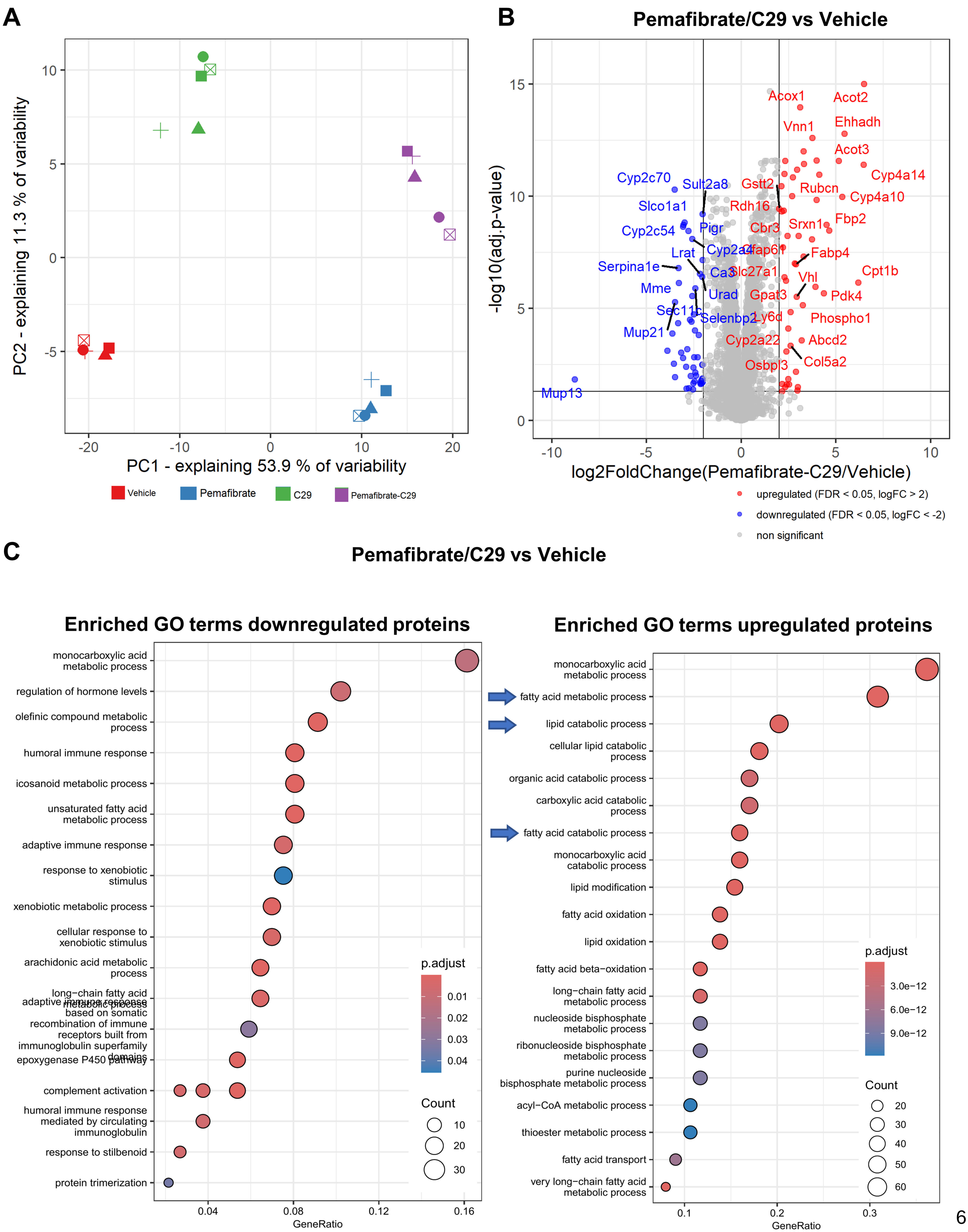

**Figure. S6**

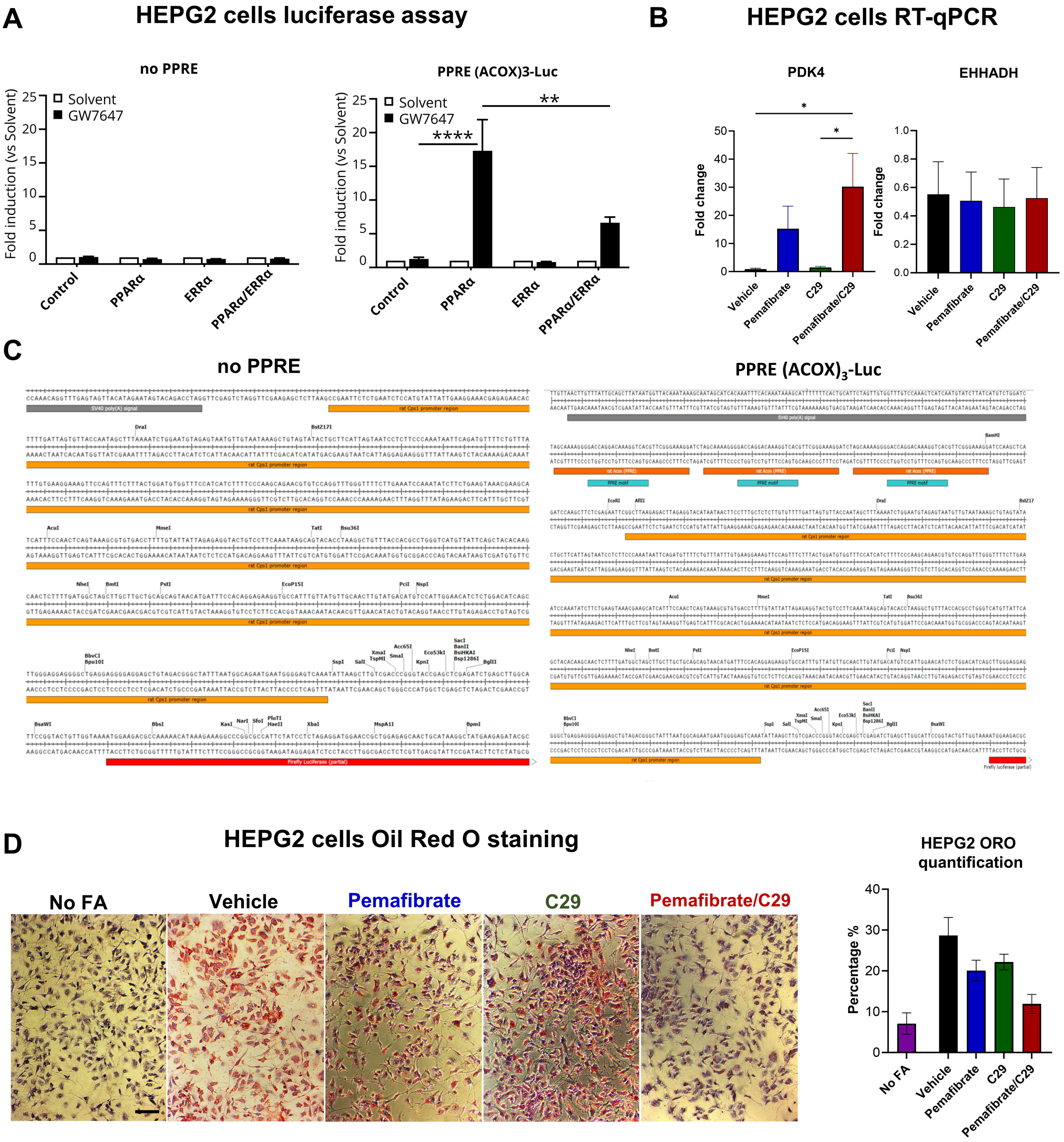
A
HEPG2 cells luciferase assay
no PPRE
PPRE (ACOX)3-Luc
Fold induction (vs Solvent)
Solvent
GW7647
****
**
Control
PPARα
ERRα
PPARα/ERRα
B
HEPG2 cells RT-qPCR
PDK4
EHHADH
Fold change
*
Vehicle
Pemafibrate
C29
Pemafibrate/C29
C
no PPRE
PPRE (ACOX)3-Luc
D
HEPG2 cells Oil Red O staining
No FA
Vehicle
Pemafibrate
C29
Pemafibrate/C29
HEPG2 ORO quantification
Percentage %

**Figure S7**

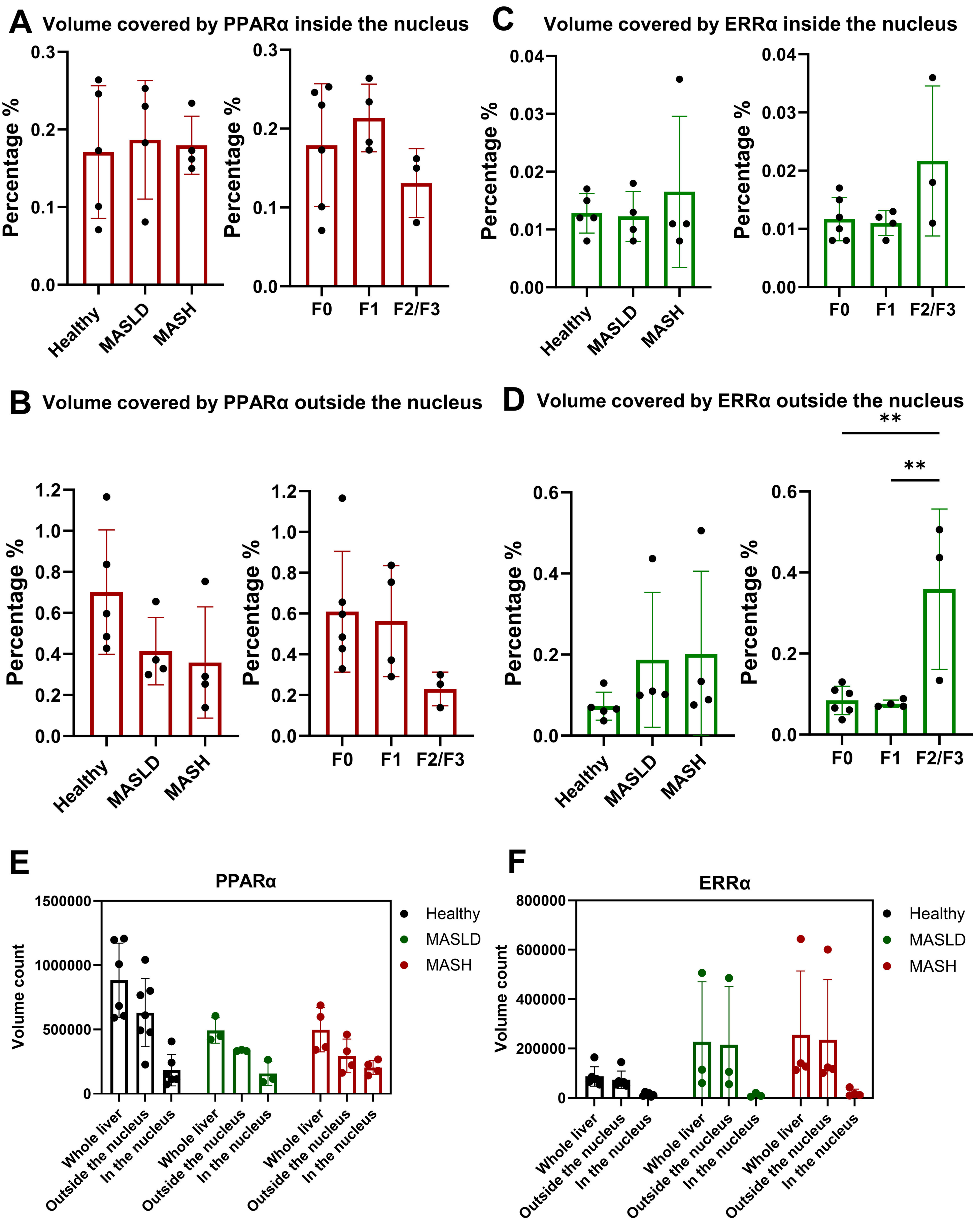
A
Volume covered by PPARα inside the nucleus
C
Volume covered by ERRα inside the nucleus
B
Volume covered by PPARα outside the nucleus
D
Volume covered by ERRα outside the nucleus
**
**
Percentage %
Healthy
MASLD
MASH
F0
F1
F2/F3
E
PPARα
Volume count
Whole liver
Outside the nucleus
In the nucleus
F
ERRα
Healthy
MASLD
MASH

#### Supplementary Figures

**Supplementary Figure 1, related to Figure 1. PPARα-ERRα ligand combination reduces steatosis and fibrosis in a long-term MASLD mice model.** 8 weeks C57Bl6 mice (n=4 mice per group) were given a western diet + fructose water for 16 weeks and treated with vehicle, Pemafibrate and/or C29 via oral gavage for 6 weeks. (A) Representative images of liver microscopic morphology taken via scanning electron microscopy of a healthy liver and (B) taken from the mice treatment group ± SD—images scale bar 1μm. The results were compared via one-way ANOVA with post-hoc testing. (*: P<0.05, **P < 0.01, ***P < 0.001, **** P < 0.0001). Abbreviations: Coll; collagen, ST; steatosis, HE; Hepatocytes, LSEC; Liver sinusoidal endothelial cells

**Supplementary Figure 2. Effect PPARα-ERRα ligand combination on gene expression in long-term diabetic Streptozotocin Western Diet mice model.** Mice were treated with vehicle, Pemafibrate and/or C29 daily via oral gavage for 6 weeks. (A) Representative liver morphology and H&E images from the mice with tumours. (B). Relative gene expression by qPCR showing tumour gene markers compared with the healthy control. (C). Representative images showing staining of H&E and tumour markers. (D-F) The relative gene expression levels of inflammation, fibrosis and lipid metabolism related genes were measured by qPCR from the STZ-WD mice model. The results were compared via one-way ANOVA with post-hoc testing. (*: P<0.05, **P < 0.01, ***P < 0.001, **** P < 0.0001).

**Supplementary Figure 3. Effect of long-term administration of PPARα-ERRα ligand combination in healthy mice.** 8-week old male mice were treated with vehicle Pemafibrate (0.1 mg/kg) and/or C29 (10 mg/kg) via oral gavage for 6 weeks. (A) C57Bl6 were given normal chow and treatments for six weeks. (B) Weekly weight measurements for 6 weeks and final weight, liver and spleen to body weight ratio. (C) Representative images of liver macroscopic morphology, H&E staining, and Sirius red staining (10 per sample at 20X). (D) Serum from mice measured for Alanine transaminase (ALT), glucose, cholesterol and triglyceride levels. (E) The relative gene expression levels of PPARα target genes were measured by qPCR from the healthy mice. All data are presented as means ± SD—images scale bar 200μm. Results were expressed as mean and were compared by one-way ANOVA with post hoc testing. (*: $p<0.05$, **$P < 0.01$, ***$P < 0.001$, **** $P < 0.0001$).

**Supplementary Figure 4, related to Figure 4. The effect of PPARα-ERRα ligand combination on gene expression in short-term MASLD mice**. 8-week-old C57BL/6J mice were fed a western diet + 10% fructose water for 14 days, and treatment was given for 7 days. Post-sacrifice, RNA was isolated from livers and subjected to RNA-sequencing. (A) Principal Component Analysis (PCA) depicting different treatment groups' variance and clustering (n=4-5 per group). (B) Volcano plot with up- and downregulated genes of pemafibrate/C9 vs vehicle filtered by $p_{adj}<0.05$ and abs(log2FC)>=1 depicted as red dots. (C) KEGG pathway analysis of pemafibrate/C9 vs vehicle-regulated genes. (D,E) RNA-sequencing-derived count plots of selected gene targets and accompanying qPCR validations.

**Supplementary Figure 5, related to Figure 4. The effect of PPARα-ERRα ligand combination on protein expression in short-term MASLD mice**. 8-week-old C57BL/6J mice were fed a western diet + 10% fructose water for 14 days, and treatment was given for 7 days. Post-sacrifice, protein was isolated from livers, further processed and subjected to MS-based shotgun proteomics. (B) Volcano plot with up- and downregulated proteins filtered by $p_{adj}<0.05$ and abs(log2FC)>=2 depicted as red or blue dots. (C) GO biological processes of proteins differentially upregulated (left) or downregulated (right) by pemafibrate/C9 vs vehicle.

**Supplementary Figure 6 ERRα blunts PPARα transcriptional activity in hepatocytes.** (A) HEPG2 cells were transfected with a CPS-luciferase reporter plasmid (negative control, no PPRE) (left panel) or a CPS-Luc variant containing three PPRE elements of the rat acyl-coenzyme A oxidase (ACOX) promoter and with different plasmids expressing ERRα and/or PPARα or control plasmid. Cells were stimulated with GW (0.5μM) or solvent control for 24h. Promoter activities were normalised versus solvent (fold induction) (N=3). (B) After overnight starvation, HEPG2 cells were stimulated with pemafibrate and/or C29 with a serum-containing medium for 24 hours. The relative gene expression level was measured (N=3). (C) Parts of the sequence of the luciferase reporter constructs. (left) The control-luciferase reporter construct contains the Cps1 promoter region, but no PPRE elements. (right) The $(ACOX)_3$-Luc reporter contains three PPRE elements of ACOX1, which precede the Cps1 promoter region. (D) HEPG2 cells were subjected to lipid stress induced by oleic acid and palmitic acid in a 2:1 (0.5-0.25mM) ratio, treated with pemafibrate and/or C29 for 24h and subjected to Oil Red O staining (ORO) (N=1). Quantitative image analysis (3 images per treatment) was performed with Fiji. All data are presented as means ± SD—images scale bar 100µm. Where applicable, results were compared via one-way ANOVA with post-hoc testing (*: P<0.05, **P<0.01, ***P<0.001, **** P<0.0001).

**Supplementary Figure 7, related to Figure 6. Expression of PPARα-ERRα receptor in healthy, MASLD and MASH patients.** Liver biopsies from Ghent University Hospital were processed for immunostaining from healthy, MASLD and MASH patients. 6 images (approximately 60 images per Z-stack) were taken per liver section with an airy scan LSM 880 super-resolution microscope, and the endogenous protein volumes (3D images) were quantified using Volocity. (A,B) Quantification results of the volume of PPARα signal inside and outside the nucleus respectively. (C,D) Quantification results of volume of ERRα inside and outside the nucleus, respectively. (E,F). Raw volume count of PPARα-ERRα. All data are presented as means ± SD—images scale bar 10µm. The results were compared via one-way ANOVA with post-hoc testing. (*: P<0.05, **P < 0.01, ***P < 0.001, **** P < 0.0001).

#### Supplementary Tables

##### Supplementary Table 1. Mice model's summary used in our study

| **2-weeks model WD + FW** | **16-weeks model WD +FW** | **12-weeks model STZ-WD** | **Healthy control 12-week model** |
|---|---|---|---|
| Treatment duration: **7 days** | Treatment duration: **6 weeks** | Treatment duration: **6 weeks** | Treatment duration: **6 weeks** |
| Vehicle | Vehicle | Vehicle | Vehicle |
| Pemafibrate (0.1 mg/kg) | Pemafibrate (0.1 mg/kg) | Pemafibrate (0.1 mg/kg) | Pemafibrate (0.1 mg/kg) |
| C29 (30 mg/kg) | C29 (30 mg/kg) | C29 (10 mg/kg) | C29 (10 mg/kg) |
| Pemafibrate/C29 (0.1 mg/kg and 30 mg/kg) | Pemafibrate/C29 (0.1 mg/kg and 30 mg/kg) | Pemafibrate/C29 (0.1 mg/kg and 10 mg/kg) | Pemafibrate/C29 (0.1 mg/kg and 10 mg/kg) |

##### Supplementary Table 2. Patient characteristics

| **Baseline characteristic** | **Control (n = 5)** | **MASLD (n=8)** | **P value** |
|---|---|---|---|
| Age, y | 38 (32-62) | 45 (42-58) | 0.435 |
| Sex, female/male | 5/0 | 5/3 | 0.231 |
| Biometry | | | |
| Weight, kg | 92.8 (79.4-131.3) | 106.5 (91.5-122.3) | 0.476 |
| BMI, m/kg² | 32.0 (28.1-46.8) | 38.6 (35.9-44.0) | 0.171 |
| Lab results | | | |
| ALT, U/L | 24 (21-58) | 44 (23-65) | 0.432 |
| AST, U/L | 24 (18-35) | 34 (18-46) | 0.530 |
| Platelets, $10^3$/µl | 264 (230-339) | 290 (242-332) | 0.876 |
| Glucose, mg/dL | 91 (78-100) | 95 (84-105) | 0.905 |
| Albumin, g/L | 46.5 (44.2-48.0) | 44.4 (41.0-44.4) | 0.857 |
| Histology | | | |
| Steatosis, grade | 0 (0-0) | 2 (1-2) | **0.002** |
| Inflammation, grade | 0 (0-1) | 1 (0-1) | 0.435 |
| Ballooning, grade | 0 (0-0) | 1 (0-2) | 0.065 |
| NAS score | 0 (0-1) | 4 (2-5) | **0.003** |
| Fibrosis, stage | 0 (0-1) | 1 (0-3) | 0.065 |

#### Supplementary Table 3. Genes and primers used for real-time qPCR on mouse samples

| NCBI Gene ID | Gene | Abbreviation | Forward primer | Reverse primer | Function/ Marker |
|---|---|---|---|---|---|
| 14433 | Glyceraldehyde-3-phosphate dehydrogenase | Gapdh | CATGGCCTTCCGTGTTCCTA | GCGGCACGTCAGATCCA | Housekeeping gene |
| 15251 | Hydroxymethylbilane synthase | Hmbs | AAGGGCTTTTCTGAGGCACC | AGTTGCCCATCTTTCATCACTG | Housekeeping gene |
| 15452 | Hypoxanthine-guanine phosphoribosyltransferase | Hprt1 | GTTAAGCAGTACAGCCCCAAA | AGGGCATATCCAACAACAAACTT | Housekeeping gene |
| 20661 | Succinate dehydrogenase complex, subunit A | Sdha | CTTGAATGAGGCTGACTGTG | ATCACATAAGCTGGTCCTGT | Housekeeping gene |
| 56636 | Fibroblast growth factor 21 | Fgf21 | CAGGGAGGATGGAACAGTGGTA | TGACACCCAGGATTTGAATGAC | Glucose metabolism |
| 14380 | Glucose 6 phosphatase dehydrogenase | G6pd | CGACTCGCTATCTCCAAGTGA | GTTGAACCAGTCTCCGACCA | Glucose metabolism |
| 12919 | Perilipin-2 | Plin2 | CTTGTGTCCTCCGCTTATGTC | GCAGAGGTCACGGTCTTCAC | Fat droplet protein |
| 18534 | Pyruvate dehydrogenase kinase 4 | Pdk4 | AGAGCCTGATGGATTTGGTG | TCCACTGTGCAGGTGTCTTT | Fatty acid oxidation |
| 14070 | Liver fatty acid binding protein | Fabp1 | CCAGGAGAACTTTGAGCCATTC | TGTCCTTCCCTTTCTGGATGT | Lipid metabolism |
| 12491 | Cluster of differentiation 36 | Cd36 | GCCAAGCTATTGCGACATGA | GAAAAGAATCTCAATGTCCGAGACT | Fatty acid transporter |
| 109652 | Acetyl-Coenzyme A acyltransferase 1B | Acaa1b | CAGGACGTGAAGCTAAAGCCT | CTCCGAAGTTATCCCCATAGGAA | Fatty acid oxidation |
| 13516 | Enoyl-CoA hydratase and 3-hydroxyacyl CoA dehydrogenase | Ehhadh | ATGGCTGAGTATCTGAGGCTG | GGTCCAAACTAGCTTTCTGGAG | Fatty acid oxidation |
| 14104 | Fatty acid synthase | Fasn | GGAGGTGGTGATAGCCGGTAT | TGGGTAATCCATAGAGCCCAG | Lipid metabolism |
| 16956 | Lipoprotein lipase | Lpl | GGGAGTTTGGCTCCAGAGTTT | TGTGTCTTCAGGGGTCCTTAG | Lipid metabolism |
| 12842 | Collagen, type I, alpha 1 | Col1a1 | AAAGGTGCTGATGGTTCTCC | GGGACCGGGAGGACCACTGG | Fibrosis |
| 21803 | Transforming growth factor-beta | Tgf-β | ACCGGCCCTTCCTGCTCCTC | GCCGCACACAGCAGTTCTTC | Fibrosis |
| 21857 | TIMP metallopeptidase inhibitor 1 | Timp1 | CTTGGTTCCCTGGCGTACTC | ACCTGATCCGTCCACAAACAG | Fibrosis |
| 20296 | Chemokine (C-C motif) ligand 2 (MCP1) | Mcp1 | TTAAAAACCTGGATCGGAACCAA | GCATTAGCTTCAGATTTACGGGT | Inflammation |
| 16193 | Interleukin-6 | Il-6 | TAGTCCTTCCTACCCCAATTTCC | TTGGTCCTTAGCCACTCCTTC | Inflammation |
| 21926 | Tumor necrosis factor-alpha | Tnfα | ACGTGGAACTGGCAGAAGAG | TCACCCCGAAGTTCAGTAGA | Inflammation |
| 19013 | Peroxisome proliferator-activated receptor alpha | Pparα | AGAGCCCCATCTGTCCTCTC | ACTGGTAGTCTGCAAACCAAA | Nuclear receptor |
| 26379 | Estrogen-related receptor alpha | Errα | AGGTGGACCCTTTGCCTTTC | GGCATGGCGTACAGCTTCT | Nuclear receptor |
| 20208 | Serum amyloid A1/2 | Saa1/2 | TTGTTCACGAGGCTTTCC | TGAGCAGCATCATAGTTCC | Acute phase inflammation |

#### Supplementary Table 4. Genes and primers used for real-time qPCR on human samples

| NCBI Gene ID | Gene | Abbreviation | Forward primer | Reverse primer | Function/ Marker |
|---|---|---|---|---|---|
| 2597 | Glyceraldehyde-3-phosphate dehydrogenase | GAPDH | TGCACCACCAACTGCTTAGC | GGCATGGACTGTGGTCATGAG | Housekeeping gene |
| 3145 | Hydroxymethylbilane synthase | HMBS | GGCAATGCGGCTGCAA | GGGTACCCACGCGAATCAC | Housekeeping gene |
| 3251 | Hypoxanthine-guanine phosphoribosyltransferase | HPRT1 | TGACACTGGCAAAACAATGCA | GGTCCTTTTCACCAGCAAGCT | Housekeeping gene |
| 6389 | Succinate dehydrogenase complex, subunit A | SDHA | TGGGAACAAGAGGGCATCTG | CCACCACTGCATCAAATTCATG | Housekeeping gene |
| 1962 | Enoyl-CoA hydratase and 3-hydroxyacyl CoA dehydrogenase | EHHADH | AAACTCAGACCCGGTTGAAGA | TTGCAGAGTCTACGGGATTCT | Fatty acid oxidation |
| 5166 | Pyruvate dehydrogenase kinase 4 | PDK4 | ACAGAGCCTGATGGATTTGGTGGA | TGACTGGGTCAACTGTACAGGCAT | Fatty acid oxidation |

### Supplementary Methods

#### Cell culture

Human hepatoma (HEPG2) cells were maintained in Dulbecco's modified Eagle medium (DMEM) (Thermofisher) supplemented with 10% fetal calf serum (FCS) with 1% Antibiotic-Antimycotic (Thermofisher) and grown at 37°C under 5% $CO_2$.

##### *HEPG2 cells - reporter assay*

HEPG2 cells were transfected with a luciferase reporter containing three PPARα-responsive elements of the acyl CoA oxidase 1 (ACOX1) promoter, or a corresponding control-luciferase reporter (negative control - no PPRE), and PPARα and/or ERRα plasmids. After 24h, cells were treated with the PPARα ligand GW7647 (GW, 0.5μM) or with solvent control (0.2% DMSO) for 24h, after which cells were lysed in 1x Cell Culture Lysis Reagent (CCLR) and luminescence and β-galactosidase was measured. Ratios of luminescence versus β-galactosidase were made and fold changes versus solvent control were plotted.

##### *HEPG2 cell setup for RT-qPCR*

Before HEPG2 cell seeding, 24-well plates were coated with a collagen/PBS +/+ mixture at a final collagen concentration of 40 μg/mL; followed by 1-hour incubation at 37°C. HEPG2 cells, seeded at 100,000 cells per well, were fasted by replacing DMEM culture medium from the wells with fetal calf serum free DMEM for 24h before ligand induction. Pemafibrate and/or C29, dissolved in 100% DMSO, were diluted to final concentrations of 5 μM in DMEM supplemented with 10% FCS (the total solvent concentration was kept similar in all conditions). Following induction for 24h, cells were harvested for RNA isolation. Experiments were performed in triplicate. Human genes used is located in supplementary 5.

##### *HEPG2 cells Oil Red O staining*

HEPG2 cells (seeded at 60,000 cells per well in 24-well plates) were incubated for 24 hours at 37°C. Hereafter, DMEM culture medium was aspirated and refreshed with DMEM without fetal calf serum containing 1% BSA and a 2:1 ratio of oleic acid and palmitic acid (0.5 mM OA

– 0.25 mM PA) (Thermofisher) for 24h. Cells were then treated with Pemafibrate (5μM) and/or C29 (5μM) for 24h.

After ligand induction, cells were gently washed 2X with PBS and Incubated with 4% PFA for 30 minutes. Then, cells were gently washed twice with Milli-Q water and exposed to a 60% isopropanol solution for 5 minutes. After removing the isopropanol, cells were completely and evenly covered with the Oil Red O solution (1:3 diluted with water). The plate or dish was incubated for 10-20 minutes under continuous shaking. The Oil Red O solution was then removed and the cells were washed 2-5 times with Milli-Q water until no excess stain was visible.

For imaging, Hematoxylin was added and the cells were incubated for 30-60 seconds. After removing the Hematoxylin, cells were washed 2-5 times as needed with Milli-Q $H_2O$. During microscopy, cells were kept covered with $dH_2O$ to ensure the lipid droplets appeared red and nuclei blue. 3 images were taken per well and quantified via Fiji (Image J). Oil red O staining with quantitative image analysis was performed in Fiji (Image J, version 2.1.0)[1].

##### Mice tissue sampling

Mice were anaesthetised via intraperitoneal injection of ketamine (60 mg/kg; Dechra Veterinary Products, Lille, Belgium) and xylazine (6 mg/kg; Kela, Sint-Niklaas, Belgium) and euthanised by cervical dislocation. The liver, spleen and adipose tissue were weighed. Two pieces of the liver were isolated; one was fixed in 4% phosphate-buffered formaldehyde solution (Klinipath, Olen, Belgium) for histological analysis, and one was incubated in RNA later (Ambion, Thermo Fisher Scientific), snap frozen in liquid nitrogen and stored at –80°C until further processing for RT-PCR.

For electron microscopy, mice samples from long-term western diet + fructose water MASLD mice model were subjected to partial hepatectomy and perfused-fixation via the left liver lobe, cut in pieces of approximately 1mm$^3$ and further fixed in 2.5% glutaraldehyde (Electron microscopy sciences, Brussels) and 4% formaldehyde in PHEM buffer (w/v: 1.81% PIPES, 0.65% HEPES, 0.38% EGTA, 0.1% MgSO4, Sigma Aldrich), pH 7.

*Partial hepatectomy and hepatic portal flushing in mice for scanning electron microscopy (SEM)*

The procedure was done in a rodent operation room, with the temperature and ambient noise controlled. Anaesthesia was induced with ketamine/xylazine as indicated. A toe pinch was applied to estimate the depth of anaesthesia, and the mice were positioned in a dorsal position on a sterile drape. The abdomen was shaved and disinfected twice with chlorhexidine solution (Fisher Scientific, Belgium). The skin was lifted using forceps, and a midline incision was cut in the abdominal wall and the linea alba (xiphoid – lower abdomen incision). Care was taken not to cut through the rectus muscles to avoid excessive bleeding. The intestines were gently eviscerated, with minimal touch of the organs, solely using moistened cotton swabs. The upper abdomen and liver lobes were visualised and exposed. The venous and portal intestinal circulation was focused on the portal and splenic veins. The separate liver lobes were identified and fanned out, ensuring the medial and left (lateral) lobes were positioned cranially, with the right superior, inferior, and caudate lobe positioned caudally. A sharp dissection of the separate lobes was performed up until the porta hepatis, thus ensuring a safe and clear view of the porta hepatis at the base of each lobe. Separate ligation of the right superior, inferior, and caudate lobe was done, either ligating with Vicryl 5/0 thread or clipping using titanium medium (blue) clips. A sharp resection and completion of the partial hepatectomy were performed. Potential bleeding was controlled using Surgipro 8/0 cross stitches. The distal vena porta was punctured, and a catheter (winged intravenous catheter, 26G, 0.64 x 19 mm) was advanced over the needle up to the porta hepatis. Once stable in place, blood was drawn for separate analysis. Infradiaphragmatic clipping of both the Vena Cava Superior and aorta was done, closing off the circulation for ideal flushing. The leftover hepatic lobes were slowly flushed. To avoid excessive pressure in the hepatocytes, the Vena Splenica was transected, ensuring a steady outflow of the perfusate. The final resection of the leftover hepatic lobes for analysis was completed.

*Serum biochemical test*

Blood was drawn from the mice via retro-orbital bleeding. It was centrifuged for 10 minutes at 10,000 rpm, and serum was collected. The samples were taken to Ghent University Hospital Laboratory for serum analysis of Alanine aminotransferase (ALT), and triglyceride and glucose levels were measured (UV test at 37°C) in serum (Roche Modular pre-analytics system, Rotkreuz, Switzerland).

For the intraperitoneal glucose tolerance test (IPGTT), mice were fasted for approximately 6 hours. The fasted blood glucose levels were determined before a glucose solution (2g/kg) (Thermo Fischer Scientific, Merelbeke, Belgium) was administered by intraperitoneal (IP) injection. Subsequently, the blood glucose level was measured simultaneously with a glucose monitor for 0, 30, 60 and 120 minutes.

#### Histochemistry

*Liver histology assessment*

Conventional hematoxylin-eosin (H&E) and Sirius red stainings were performed according to established protocols[1]. Histological features were scored according to the MASH clinical research network scoring system by a researcher blinded to the treatment groups[2,3]. Sirius red area fractions were analysed and quantified (ImageJ, public domain). Fibrosis was evaluated using the MASH Clinical Research Network fibrosis staging system[2,3]. Any tumour signs observed were categorised as none, small, medium and large. A diagnosis of Metabolic dysfunction-associated steatotic liver disease (MASLD) fatty liver was made if ≥5% of hepatocytes contained macrovesicular lipid droplets, whereas the diagnosis of MASH was based on the joint presence of steatosis, hepatocyte ballooning and lobular inflammation.

*Immunohistochemistry*

Immunohistochemical analysis was performed on formalin-fixed, paraffin-embedded tissue sections to assess the expression of Glypican 3, HSP70, and Glutamine Synthetase. The sections were first deparaffinized by placing them in xylene for three changes of 5 minutes each. This was followed by rehydration through immersion in a graded series of ethanol solutions: 100% ethanol for 5 minutes, 95% ethanol for 5 minutes, 70% ethanol for 5 minutes, and finally distilled water for 5 minutes.

Antigen retrieval was conducted using Cell Conditioning 1 (CC1) solution at 95°C. The incubation times varied depending on the target protein: Glypican 3 and Glutamine Synthetase were incubated for 36 minutes, while HSP70 was incubated for 32 minutes. To block non-specific binding sites, sections were incubated with 5% normal serum for 10 minutes at room temperature.

The primary antibody incubation was carried out at 37°C. For Glypican 3, the Roche Diagnostics antibody (Ref.: 06483186001), provided ready to use, was applied for 32 minutes. For HSP70, the Santa Cruz Biotechnology antibody (Ref.: sc-24) was diluted 1:100 and incubated for 24 minutes. Glutamine Synthetase detection utilized the Roche Diagnostics antibody (Ref.: 07107757001), also ready to use, and was incubated for 12 minutes.

Following the primary antibody application, a suitable secondary antibody conjugated to horseradish peroxidase (HRP) was applied to the sections for 30 minutes at room temperature. Visualization of the target proteins was achieved using specific detection kits: the UltraView Universal DAB Detection Kit for Glypican 3 and Glutamine Synthetase, and the OptiView DAB IHC Detection Kit for HSP70. The DAB substrate-chromogen solution was applied, and the slides were incubated for 5-10 minutes until the desired staining intensity was reached. The sections were then rinsed in distilled water.

Counterstaining was performed using hematoxylin for 1-2 minutes, followed by rinsing under running tap water for 5 minutes to enhance the blue hue of the nuclei. The slides were then dehydrated through sequential immersion in 70% ethanol, 95% ethanol, and 100% ethanol for 5 minutes each, and cleared in xylene for two changes of 5 minutes each. Finally, the sections were mounted with a coverslip using a suitable mounting medium.

The stained sections were examined under a light microscope to evaluate the expression and localization of Glypican 3, HSP70, and Glutamine Synthetase.

##### *Oil red O tissue staining*

The tissue was first incubated in a 4% PFA (VWR International BV, Amsterdam, Netherlands) solution, which is commonly used for fixation before embedding with paraffin, for one hour at 4°C while shaking; a shaker was placed in the fridge for this purpose. Subsequently, the tissue was washed with PBS. Overnight, the tissue was incubated in a 30% sucrose solution dissolved in PBS at 4°C. The sucrose solution was prepared in a water bath for faster dissolving. For preparation, a layer of OCT (Optimal Cutting Temperature compound) was placed in the cryomold, followed by the tissue, and then another layer of OCT (VWR International BV). The setup was then frozen on dry ice and subsequently stored at -20 °C until sectioning.

The oil red O (ORO) stock solution (Sigma-Aldrich, Belgium) was prepared by adding 2.5g of ORO to 400 ml of 99% isopropyl alcohol (Sigma-Aldrich) and stirring magnetically for 2 hours at room temperature (RT; 20–25 °C). 1.5 parts of the ORO stock solution were mixed with one part of distilled water for the ORO working solution. The solution was allowed to stand for 5 minutes at 4°C, during which it thickened, then filtered through a 45-μm filter to remove precipitates and used within 6 hours. A Leica CM 1950 (Leica Microsystems Belgium BV) was used to make 12-μm-thick liver sections, and the microtome temperature was maintained at -23 degrees Celsius. To prevent detachment during the ORO staining procedure, the sections were allowed to dry for 10 minutes at RT before being frozen. The ORO staining procedure involved air drying the frozen sections on slides for a minimum of 30 minutes and then fixing them in 4% PFA for 30 minutes. The sections were quickly dipped once in 60% isopropanol (Propanolol-4), stained in working ORO solution for 3 minutes, and then quickly dipped again in 60% isopropanol and once in deionised water. Sections were counterstained with Mayer’s Hematoxylin (Sigma-Aldrich, Belgium) for 30 seconds, followed by 10 dips in deionised water. Finally, coverslips were placed on the sample with an aqueous mounting gel (VWR, Belgium).

It was crucial not to press on the coverslips to remove air bubbles as the lipids might move to the coverslip immediately after DI water to avoid air drying.

*Scanning electron microscopy (SEM)*

SEM was performed according to the protocol[1]. 5 random overview images were taken per liver sample imaged at 1000X magnification. They were imaged using Zeiss Crossbeam 540 FIB-SEM (Carl Zeiss, Germany). Steatosis (holes present) was quantified via Fiji, Image J and fibrosis were categorised as none, small, medium and large.

*PPARα-ERRα double immunofluorescence staining*

Sequential immunostaining was performed on 4-micrometre thick Formalin-Fixed Paraffin-Embedded (FFPE) murine liver sections with a microtome (Thermo HM340E, Thermofischer Scientific). The protocol was initiated with an overnight incubation of the tissue sections at 60°C to dissolve and remove the paraffin embedding medium effectively. Deparaffinisation was meticulously performed through a series of washes: thrice in Xylene for 5 minutes each to clear the paraffin, followed by a descending alcohol series (99%, 96%, and 70% ethanol) to rehydrate the tissues, and finally, a rinse in distilled water. Antigen retrieval, a critical step for unmasking epitopes, was conducted by immersing the tissue sections in 1x citrate buffer pre-warmed in a water bath. The sections were treated at 95°C for 30 minutes in this buffer, followed by a gradual cooling period of 30 minutes at room temperature. After marking the sections with a Dako pen to delineate the area of antibody application, they were washed in PBST (0.1% Tween 20 in PBS) to remove any remaining fixative and to permeabilise the tissues. To reduce non-specific binding, a blocking solution comprising 5% Donkey serum and 5% Goat Serum (VWR, Belgium) in PBST was applied to the sections for one hour. Notably, the protocol emphasized the omission of a washing step post-blocking to maintain the integrity of the blocking layer. The primary antibodies were then diluted in the same blocking buffer and applied to the sections. PPARα (Anticorps PPARα (H-2): CAT: sc-398394, Santa Cruz Biotechnology) and (ERRα catalogue number ab76228, Abcam) staining was performed at a dilution of 1:100. Post-primary antibody incubation, the sections were thoroughly washed in PBST to remove any unbound antibodies. The secondary antibodies, diluted in blocking buffer, were then applied for one hour at room temperature in a dark environment to prevent photobleaching. After secondary antibody (1:800 dilution) application with Alexa flour 488 goat anti-rabbit (Invitrogen, Invitrogen: A32731), and Alexa flour 568 donkey anti-mouse (Invitrogen, CAT: A10037) sections were washed extensively with PBS to remove any excess antibodies. The sections were then treated with DAPI for nuclear counterstaining and subsequently washed in water. Vectashield (H-1000) was carefully applied for mounting, and the cover glasses were sealed with a mounting medium. The prepared slides were then stored in a 4ºC cold room until microscopic analysis and were imaged the following day.

*Confocal imaging and 3D velocity analysis*

Confocal images (8-bit) were captured with an LSM880 confocal microscope equipped with an Airyscan detector (Zeiss, Jena, Germany). 6 images (approximately 60 Z-stack per image) were taken in super-resolution, FAST mode by using a Plan-Apochromat 40x/1.2 oil objective. AF 488 was excited using the 488 nm line of an Ar laser (5%), and emission was captured between 495 and 550 nm. AF 568 was excited by a diode laser at 579 and 603 nm. Z-sections were made every 100 nm. Images were calculated through pixel reassignment and Wiener filtering by using the built-in "Airyscan Processing" command in the Zen software. Segmentation and overlap were done in 3D using Volocity software (Quorum Technologies). PPARα and ERRα positive objects (groups of pixels with intensity values above a pre-defined threshold) were segmented and calculated for these objects.

##### RNA extraction and RT-qPCR

RNA was extracted from 20 mg mouse liver tissue, preserved in RNA later (Invitrogen), using the RNeasy plus mini kit (Qiagen), according to the manufacturer's protocol. The RNA quality was evaluated by spectrophotometry, calculating the A260/A280 and A260/230 ratio's. cDNA synthesis was performed starting from 1µg RNA, using the SensiFAST cDNA synthesis kit (Bioline, London, UK). cDNA served as a template for the QPCR reaction. cDNA was added to a 384-well plate with specific primers (Biolegio, Nijmegen, The Netherlands) (Supplementary Table 3) and Sensimix SYBR No-ROX Mastermix (Bioline). Samples were run and analysed on the Lightcycler 480 II (Roche). PCR reactions using water instead of a template showed no amplification. Technical duplicates were used and Cq values were calculated with the second derivative maximum method. Average Cq values were normalised to the Cq of stable reference genes, according to GeNorm analysis in qBase+(Biogazelle, Ghent, Belgium).

RNA was isolated from cells using the RNeasy Micro Kit (Qiagen, Cat. N°: 79654, and 74106), according to the manufacturer's protocol (Invitrogen). cDNA was isolated using a cDNA synthesis kit from Biotech Rabbit (Cat. N° = BR0400404), and served as a template for the QPCR reaction (from here, a similar workflow as above). Gapdh, Hmbs, Hprt1, Sdha, or Actb served as reference genes, with primers in Supplementary Table 4.

#### RNA sequencing

Total RNA was extracted utilizing the RNeasy mini kit (Qiagen), and the RNA-seq library was constructed with the Illumina TruSeq stranded mRNA library kit. Following the library prep, the samples were sequenced on a Illumina NovaSeq 6000 instrument (VIB Nucleomics core), following these parameters: Illumina NovaSeq 6000 v1.5 sequencing kit 100 cycles single-end reads (101-10-10-0), 1% PhiX. An average of 20 million reads/sample was achieved. Sequencing reads were subjected to the nf-core pipeline (version 3.6)[4] to generate count data. This encompassed initial quality check with FastQC (version 0.11.9) and trimming of reads with Trim-Galore (version 0.6.7) to eliminate low-quality ends (phred score<30) and adapters.

A subsequent quality assessment of the trimmed data was performed. The reads were then aligned to the mouse genome GRCm38 using STAR (version 2.6.1d) and converted into count data using SALMON (version 1.5.2). Differential gene expression analysis, on genes with counts greater than one, was conducted using the DESeq2 R package (version 1.38.3). Pairwise comparisons between treated samples were determined at a significance level of $\alpha=0.05$, corresponding to the adjusted p-value (FDR) cutoff ($p_{adj}$). The $p_{adj}$ (cutoff: < 0.05) and log2FC (cutoff: > 1 or <1) for each pairwise comparison. Heatmaps were generated using pheatmap (version 1.0.12), clustering the differentially expressed genes for several pairwise comparisons based on log2FC. Venn diagrams were generated using ggvenn (version 0.1.10). Volcano plots were generated using EnhancedVolcano (version 1.16.0) to display the $p_{adj}$ (log10 scale) as a function of the log2FC for all genes and each pairwise comparison of interest. Herein, the gene names are indicated for the top 10 genes having the largest -log10($p_{adj}$). Functional annotations and KEGG pathway analyses of differentially expressed genes were done using gprofiler2 (version 0.2.2).

**STRING analysis**

Pairwise contrasts (e.g. PemaC29 vs vehicle) from the DESeq2-based differential gene expression analysis were fed to STRING (version 11.5, available at https://string-db.org) to perform functional enrichment analysis and to identify functional protein association networks. For functional enrichment analysis, we used FDR stringency medium (5%) and sorted on enrichment score. We focused on Gene Ontology (GO) biological processes. Specifically, we selected the "inflammatory response" category (GO:0006954) to explore pathways related to inflammation. Further enrichment analysis highlighted processes associated with "acute phase response" (GO:0002526) and "acute inflammatory response" (GO:0006953). Functional protein association networks were build using the gene targets behind the terms "inflammatory response" and "acute phase response". The analysis was performed using the "full STRING network" setting, with a minimum required interaction score set to "medium confidence" and a medium FDR stringency (5%).

#### Mass Spectrometry-based Shotgun Proteomics

*LC-MS/MS analysis*

Peptides were re-dissolved in 20 µl loading solvent A (0.1% trifluoroacetic acid in water/acetonitrile (ACN) (99.5:0.5, v/v)) of which 1 µl was injected for LC-MS/MS analysis on an Ultimate 3000 ProFlow nanoLC system in-line connected to a Orbitrap Exploris 240 mass spectrometer (Thermo). Trapping was performed at 20 µl/min for 2 min in loading solvent A on a 5 mm trapping column (Thermo scientific, 300 µm internal diameter (I.D.), 5 µm beads). The peptides were separated on a 250 mm Aurora Ultimate, 1.7µm C18, 75 µm inner diameter (Ionopticks) kept at a constant temperature of 45°C. Peptides were eluted by a gradient reaching 26.4% MS solvent B (0.1% FA in acetonitrile) after 80 min, 44% MS solvent B at 95 min, 56% MS solvent B at 100 min followed by a 5-minutes wash at 56% MS solvent B and re-equilibration with MS solvent A (0.1% FA in water). The flow rate was set to 250 nl/min.

The mass spectrometer was operated in data-independent mode, automatically switching between MS and MS/MS acquisition. Full-scan MS spectra ranging from 400-900 m/z with a normalized target value of 300%, a maximum fill time of 25 ms and a resolution at of 60,000 were followed by 30 quadrupole isolations with a precursor isolation width of 10 m/z for HCD fragmentation at an NCE of 30% after filling the trap at a normalized target value of 2000% for maximum injection time of 45 ms. MS2 spectra were acquired at a resolution of 15,000 with a scan range of 200-1800 m/z in the Orbitrap analyser without multiplexing. The isolation intervals were set from 400 – 900 m/z with a width of 10 m/z using window placement optimization.

The polydimethylcyclosiloxane background ion at 445.120028 Da was used for internal calibration (lock mass) and QCloud has been used to control instrument longitudinal performance during the project[5,6].

*Data-analysis*

LC-MS/MS runs of all samples were searched together using the DiaNN algorithm (version 1.8.1), library free. Spectra were searched against the human protein sequences in the Swiss-Prot database (database release version of 2022_01), containing 20,588 sequences (www.uniprot.org) and the common contaminants database protein sequences[7]. Enzyme specificity was set as C-terminal to arginine and lysine, also allowing cleavage at proline bonds with a maximum of two missed cleavages. Variable modifications were set to oxidation of methionine residues and acetylation of protein N termini. Mainly default settings were used, except for the addition of a 400-1000 m/z precursor mass range filter and MS1 and MS2 mass tolerance was set to 15 and 20 ppm respectively. Further data analysis of the shotgun results was performed with an in-house script in the R programming language, version 4.2.2. Protein expression matrices were prepared as follows: the DIA-NN main report output table was filtered at a precursor and protein library q-value cut-off of 1 % and only proteins identified by at least one proteotypic peptide were retained. After pivoting into a wide format, iBAQ intensity columns were then added to the matrix using the DIAgui's R package get_IBAQ function[8]. PG.MaxLFQ intensities were log2 transformed and replicate samples were grouped. Proteins with less than three valid values in at least one group were removed and missing values were imputed from a normal distribution centered around the detection limit (package DEP[9]) leading to a list of 5,980 quantified proteins in the experiment, used for further data analysis. To compare protein abundance between pairs of sample groups combination treatment vs single treatment groups, statistical testing for differences between two group means was performed, using the package limma[10]. Statistical significance for differential regulation was set to a false discovery rate (FDR) of <0.05 and |log2FC| >=2). Z-scored LFQ intensities from significantly regulated proteins were plotted in a heatmap after non-supervised hierarchical clustering.

#### RNA-seq and Proteomics integration

Following differential expression analysis, proteomics and RNA-seq data were integrated using R version 4.3.3, biomaRt package 2.58.2 and mouse GRCm39 genome Ensembl release 112. We matched proteomics data to genes using the first UniProt ID from a protein group, relying on the UniProt ID to Ensembl gene ID relationship from Biomart, prioritizing genes from the primary assembly and transcript isoforms corresponding to SwissProt entries over TrEMBL. The remaining unmatched proteins were either linked to RNA-seq data by gene name or removed.We extracted significant differentially expressed (DE) proteins and transcripts from each statistical comparison using an adjusted p-value of 0.05 and log2 fold change (log2FC) absolute value of 1 as thresholds.
